# Current Reporting Practices in Human Neuroscience Research

**DOI:** 10.1101/2024.09.28.615619

**Authors:** Arianna M. Gard, Deena Shariq, Alison A. Albrecht, Alaina Lurie, Hyung Cho Kim, Colter Mitchell, Luke W. Hyde

## Abstract

Concerns for the replicability, reliability, and generalizability of human neuroimaging research have led to intense debates over sample size and open science practices, with more recent attention on the contributions of sampling and recruitment practices. Key to understanding the state of neuroscience research is an assessment of reporting practices that influence replicability, reliability, and generalizability. In this structured review, we evaluated reporting practice across three domains: (1) demographic (e.g., reporting participant race-ethnicity, age, any measure of socioeconomic position), (2) methodological (e.g., reporting recruitment methods, inclusion and exclusion criteria, why participants were excluded from analyses), and (3) open science and generalizability (e.g., analyses were preregistered, target population was stated). Included were 919 published MRI and fMRI studies from 2019 in nine top-ranked journals (N = 3,856 records screened). Reporting across domains was infrequent, with participant racial or ethnic identity (14.8%), reasons for missing imaging data (31.2%), and identification of a target population (19.4%) being particularly low/underreported. Reporting likelihood varied by study characteristics (e.g., participant age group) and was correlated across domains. The median sample size of studies was 55 participants. Study sample size, reporting frequency, was positively associated with two-year citation counts, providing some evidence that the complete reporting of demographic characteristics, methodological decisions, and open science and generalizability practices may not be as valued as study sample size. Recommendations for structural interventions at the journal level are proposed.

## 1. Introduction

Over the last several decades, structural and functional neuroimaging have become widespread tools to study thoughts, emotions, behavior, and health. The human brain is a topic of public fascination, with neuroscience research impacting major legal decisions (e.g., Roper v. Simmons, [2005]) and the subject of global scientific initiatives (e.g., the National Institutes of Health BRAIN initiative; Yuste & Bargmann, 2017). Brain science further informs other fields (e.g., philosophy; Bennett & Hacker, [2022]), impacts what we think makes us human (Greene, 2014), and even which behaviors we believe are “normal” versus “abnormal” (Gazzaniga, 2005). Moreover, neuroscience has the potential to inform the diagnosis, prognosis, and treatment of many diseases (Volkow & Boyle, 2018).

However, for any scientific research to be useful to practitioners, policymakers, or the public, study findings must be reliable (i.e., the construct can be reproduced from repeated measurements), replicable (i.e., analysts can reproduce the same results given the same data and methodological plan), and externally valid (i.e., study findings generalize to a broader population). Reliability, replicability, and validity are the bedrock of the scientific method and have been the subject of debate for as long as science itself. Neuroscience research is no different, but the role of brain phenotypes in supporting individual differences and the nature of how we collect and prepare brain data for analyses makes maintaining reliability (Elliott et al., 2020), replicability (e.g., Botvinik-Nezer et al., 2020), and external validity (Falk et al., 2013; Gard et al., 2023) particularly challenging. t

Most discussions on how to improve the utility of human neuroimaging research center on the links between sample size, statistical power, and reliability (Turner et al., 2018) and the adoption of open-science practices that promote reproducibility, such as posting reproducible code and pre-registering hypotheses (Nichols et al., 2017). Empirical investigations have revealed several methodological decisions that influence reliability, replicability, and generalizability. For example, test-retest reliability is larger using multivariate versus univariate statistical approaches (Marek et al., 2022), with more data (Elliott et al., 2020), and when examining within-person changes in brain-behavioral development (Flournoy et al., 2024; Gratton et al., 2022). Such empirical investigations are critical. To more completely gauge the state of neuroscience research, however, we need to understand the decisions that researchers make (and report) in the production of this knowledge – from the strategies to recruit participants to the data filtering that leads to the ultimate analytic sample size. This structured review evaluates reporting practices in nearly 1000 published MRI and fMRI neuroimaging studies across three comprehensive domains: demographics (e.g., age, sex, racial-ethnic identity, sample size), methods (e.g., recruitment method, quality control), and open science and generalizability.

### 1.1. Sample Sociodemographic Composition and Population Generalizability

Following longstanding calls to diversify sample representation in psychological research (Arnett, 2008; Henrich et al., 2010), neuroscience researchers have begun to attend to the sociodemographic composition of human neuroimaging studies as a key indicator of population generalizability (Dotson & Duarte, 2020; Falk et al., 2013; Garcini et al., 2022; Habibi et al., 2015; Kopal et al., 2023; Qu & Telzer, 2017; Rowley & Camacho, 2015; Weinberger et al., 2020). Empirical evaluations of sample composition in select journals (Dotson & Duarte, 2020) or topical areas (Qu et al., 2021), as well as perspective pieces on the state of the field (e.g., Kopal et al., 2023; Weinberger et al., 2020), indicate trends reflected broadly in scientific research: participants in neuroimaging studies are more likely to be socioeconomically-advantaged, of majority racial-ethnic and cultural groups, and/or from White-Educated-Industrialized-Rich-Democratic (WEIRD) populations (Henrich et al., 2010).

Sample diversity along intersecting sociodemographic dimensions (e.g., race-ethnicity, cultural orientation, social and economic resources, geography) has direct implications for external validity. In turn, sample composition impacts our basic science understanding of human beings and the application of precision medicine techniques to reduce disease burden and promote wellbeing. For example, when predictive models using neuroimaging data are trained in homogenous samples, the models perform worse in samples with greater sociodemographic diversity (Greene et al., 2022). Thus, although leveraging neuroimaging data for precision medicine is an exciting and cutting-edge opportunity, these advancements could risk excluding already-marginalized populations who might benefit the most from personalized medicine interventions (for an example in the field of statistical genetics, see Martin et al., 2019). In another example, Gard et al. (2023) leveraged the Adolescent Brain Cognitive Development® (ABCD) Study to examine the influence of population sampling weights on the associations between socioeconomic resources and several metrics of brain development. Although the study incorporated a sampling design intended to recruit a diverse sample of children aged 9-10-years at baseline, children in the recruited sample were significantly more socioeconomically-advantaged along some dimensions than the target population of same-aged youth (Gard et al., 2023; e.g., 42.1% of children in the sample had household incomes of > 400% of the federal poverty line, compared to 29.9% in the target population). In models that adjusted for sampling weights, which up- or down-weighted participants so that the sample demographics matched the target population, associations with household income were weaker than with caregiver education; unweighted models led to the opposite conclusion. These results demonstrate that different conclusions about the nature of SES associations with the brain may emerge as a function of sample sociodemographic composition (Gard et al., 2023).

### 1.2. Sampling and Recruitment

How we recruit and retain research participants impacts sample sociodemographic composition. Most studies in neuroscience, psychology, and psychiatry adopt convenience sampling approaches, in which participants are recruited from community settings (e.g., clinics, university participant pools, registry databases), flyers, online advertisements, and word-of-mouth (Nielsen et al., 2017). Though cost effective, convenience sampling approaches are highly subject to selection bias (Oswald et al., 2013) in the form of underrepresentation of large segments of the population to which researchers are trying to generalize (i.e., the “target population”). This is not always the case; recruitment strategies that are tailored to multiple sociodemographic groups (e.g., hiring “cultural insiders”, extensive outreach activities, including participant in study design) are successful in recruiting diverse samples (Habibi et al., 2015).

Once participants are recruited to participate in a neuroimaging study, eligibility requirements and quality control procedures further reduce the number of “usable” participant data to a final analytic sample. The process is described as the “flow of participants” by the APA (American Psychological Association, 2020) and is a key component of results sections (e.g., “Give reasons for nonparticipation at each stage”; “Consider the use of a flow diagram”) as outlined in STROBE reporting guidelines (Elm et al., 2008). Each decision with respect to eligibility/ineligibility, quality control (e.g., removing participant with low behavioral performance), and missing data (e.g., listwise deletion, imputation) influences the sociodemographic makeup of the final analytical sample. Moreover, information from each of these decisions can inform “where” in the pipeline participants are being lost and lead to methodological improvements that promote inclusivity. For example, extensive research linking movement during MRI to participant age has led to best practices in child neuroimaging studies to include a practice session with a “mock scanner” (Perlman, 2012) and software that provides real-time motion feedback (Dosenbach et al., 2017). Thus, to evaluate the degree of population generalizability in human neuroimaging studies and to develop protocols to improve recruitment and retention, we need a clear understanding of where and how participants are recruited from and the decisions that contribute to the composition of the final analytic sample.

### 1.3. Open Science and Generalizability

Even in the case where we successfully recruit and retain a sample that mirrors the sociodemographic composition of target population, some research practices can undermine reliability, replicability, and generalizability. Open-science tools including pre-study power analyses and preregistration (Nosek et al., 2018) can guard against questionable scientific practices such as p-hacking (i.e., estimating dozens of models with various specifications until statistical significance reaches an established threshold) and HARKing (i.e., hypothesizing after the results are known; Kerr [1998]). Recent discussions of low test-retest reliability of brain data and low reproducibility of study findings reiterate the importance of open science practices. For example, together with 70 independent teams of researchers, Botnivik-Nezer et al. (2020) analyzed the same fMRI dataset and found substantial variability in the results of nine a-priori hypotheses – only one hypothesis showed a high rate of statistically-significant findings. Variation was due to analytic decisions within the data cleaning and analytic processing streams (Botvinik-Nezer et al., 2020). ‘Multiverse analyses’ (Simonsohn et al., 2019), preregistration (Nosek et al., 2018), and posting raw data in open repositories such as NeuroVault (Gorgolewski et al., 2015) are all cited as paths to improving replicability.

Although not traditionally considered a component of ‘open science’, an author’s assessment of the generalizability of scientific findings is also a measure of transparency. This includes a clear description of the target population to which a study seeks to generalize (Simons et al., 2017), and a discussion of study limitations related to external validity (American Psychological Association, 2020; Elm et al., 2008). Although metascience work has examined reporting practices that contribute to reliability, replicability, and generalizability (e.g., Qu et al., 2021), the extent to which individual research papers comment on the intended target population or provide an assessment of external validity is yet unknown.

### 1.5. Empirical Evaluation of Reporting Practices

In summary, empirical investigations have identified methodological factors that contribute to the reliability, replicability, and generalizability of human neuroscience research. Missing is a comprehensive assessment of the extent to which key methodological factors are *reported* in existing published works across subfields of human neuroscience. This knowledge is crucial to understanding the state of the field, including who this research represents and the methodological decisions that may contribute to scientific generalizability. An exploration of which study characteristics predict reporting frequency would also help to identify strategies to promote greater transparency (e.g., are some study types particularly susceptible to low reporting rates). Lastly, given the strong emphasis on sample size in recent years for improving the utility of human neuroscience research (Turner et al., 2018), more research is needed to understand how pressure for larger sample sizes intersects with reporting transparency and the extent to which sample size and reporting frequency are inherently valued in the field. The current study attempts to address these knowledge gaps through a systematic evaluation of reporting practices of all MRI and fMRI studies published in 2019 in top-ranked journals in psychology, psychiatry, and neuroscience.

The first aim was to document reporting rates with regards to sample demographic features (e.g., age, sex, racial-ethnic identity, sample size), methods (e.g., recruitment method, quality control), and open science and generalizability practices (e.g., pre-registration, specification of a target population). Given the foundational role of study design in research methods training, we hypothesized that studies would report methodological features at higher rates than demographic information and open science and generalizability practices. No specific hypotheses about the exact rates of reporting were generated. The second aim was to evaluate how study characteristics were associated with reporting in all three domains, with no explicit hypotheses. Third, the associations between study characteristics and sample size were evaluated, providing a much-needed update of previous metascience investigations of human neuroimaging studies (e.g., Button et al., 2013). We hypothesized that studies with larger sample sizes would be more likely to be composed of adult-aged participants (i.e., due to lower compliance and data quality in child samples) and leverage sMRI (i.e., due to the large influence of motion artifacts in fMRI). The last aim was to evaluate to the degree to which reporting likelihood and study sample size were “valued” by the field, using citation counts as an indirect measure of value. Although we hypothesized positive associations between study sample size and reporting likelihood and citation count, no comparative hypotheses were generated.

## 2. Methods

### 2.1. Identification and selection of studies

This systematic review was conducted using PRISMA guidelines (Moher et al., 2009), and the protocol was pre-registered (https://osf.io/6tpsh/) before target articles were identified, coded, or analyzed. Training procedures and deviations from the registered protocol are described in Supplemental Methods and Supplemental Table 1.

To evaluate current methodological practices in human neuroimaging studies, we first selected journals to identify target articles published in 2019. The Web of Science Journal Citation Reports tool (Clarivate, 2024) was used to sort journals by impact factor within the categories of “Psychiatry”, “Neuroscience”, and “Neuroimaging”. To balance the scope of the review with team resources, we chose the two highest-ranked journals in each category with the following exclusion criteria: (1) journals that primarily publish review papers, (2) journals that are not indexed on PubMed or PsychInfo, and (3) journals that do not include the words “neuroimaging” or “neuroscience” in the public statement of aims and scope. We also included three specialty journals for their coverage of human neuroimaging research in cognitive, affective, and developmental domains (i.e., *Developmental Cognitive Neuroscience*, *Journal of Cognitive Neuroscience*, and *Social Cognitive and Affective Neuroscience*). In total, nine journals were included in the systematic review (Supplemental Table 2). Next, in July 2020, all articles published during 2019 in any of the nine selected journals were imported from PubMed or PsychInfo into the systematic review software DistillerSR (*DistillerSR*, 2023).

All titles and abstracts were screened by two trained research assistants, with conflicts adjudicated by the first author (Polanin et al., 2019). Exclusion criteria included: (1) the article is a review paper, commentary, systematic review, meta-analysis, case study, or methods paper/technical report, (2) the study subjects are non-human subjects (e.g., animals, cell lines), and (3) the study does not include MRI or fMRI data. Articles that included both MRI/fMRI and another imaging modality (e.g., fNIRS, EEG, MEG, PET, SPECT, TMS) were excluded.

### 2.2. Data Extraction

A coding system was developed by the first author and last two authors. Codes were designed to capture the process of conducting a neuroimaging study, from recruitment to generalizing study findings to a broader population. Categories of codes included (1) global study features (e.g., participant age, study type [observational/intervention], imaging modality, smallest and largest analytic sample size), (2) sociodemographic information (e.g., gender or sex, race-ethnicity, socioeconomic resources), (3) methods (e.g., report of recruitment procedures, inclusion/exclusion criteria, quality control procedures, missing values analyses), and (4) generalizability and open science (e.g., power analyses, pre-registration, identification of target population). The codebook was generated using reporting guidelines from the APA Style Guide 7^th^ Edition (American Psychological Association, 2020) and STROBE (Elm et al., 2008). Table 1 provides a summary of the codebook; the complete codebook can be found in the Supplementary Table 3. Throughout the training phase and during the first several months of coding, the coding manual was updated iteratively with definitions of terms (e.g., intervention study criteria) and data entry instructions (e.g., round numbers to the nearest percent).

**Table 1.**
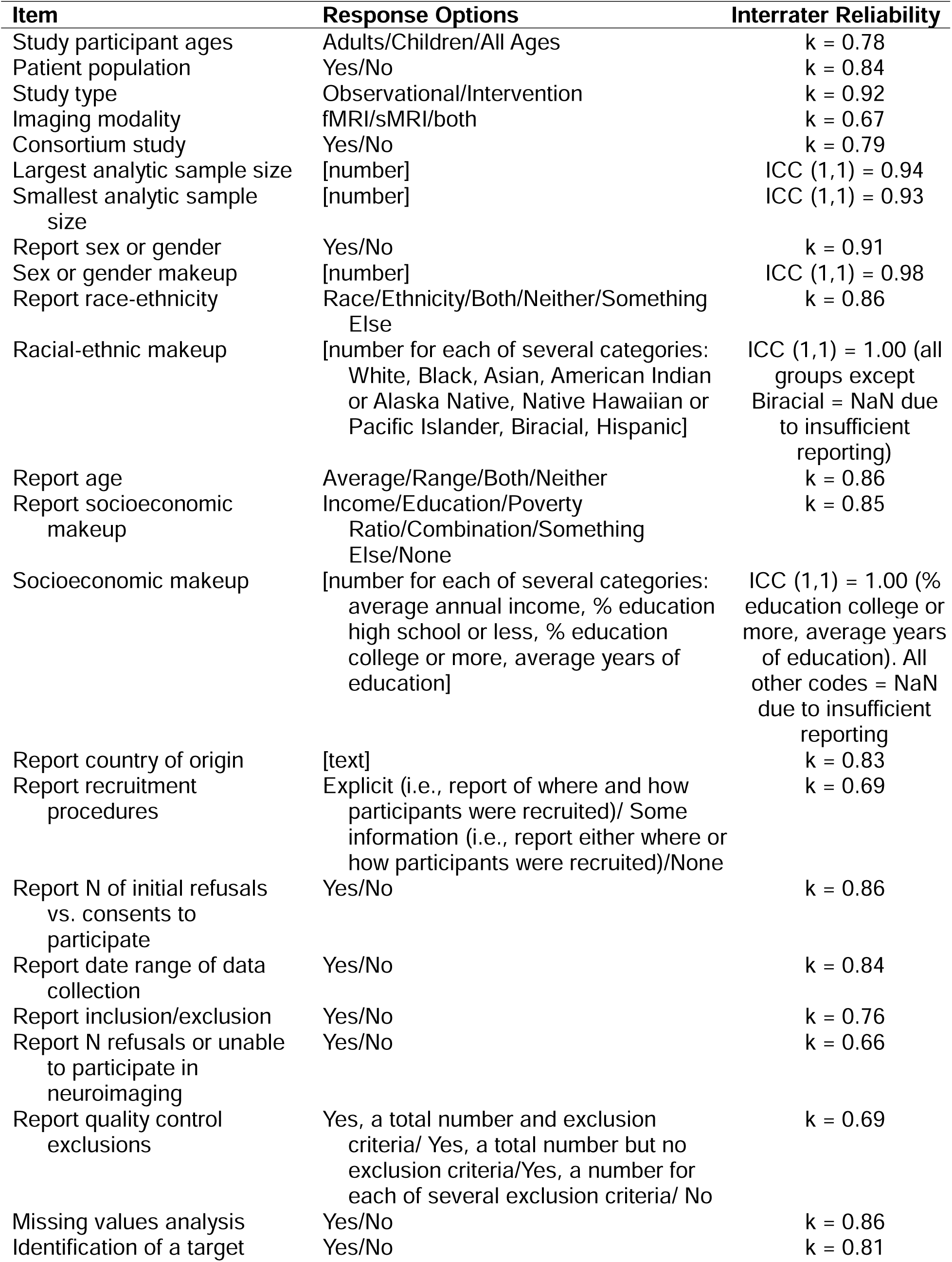
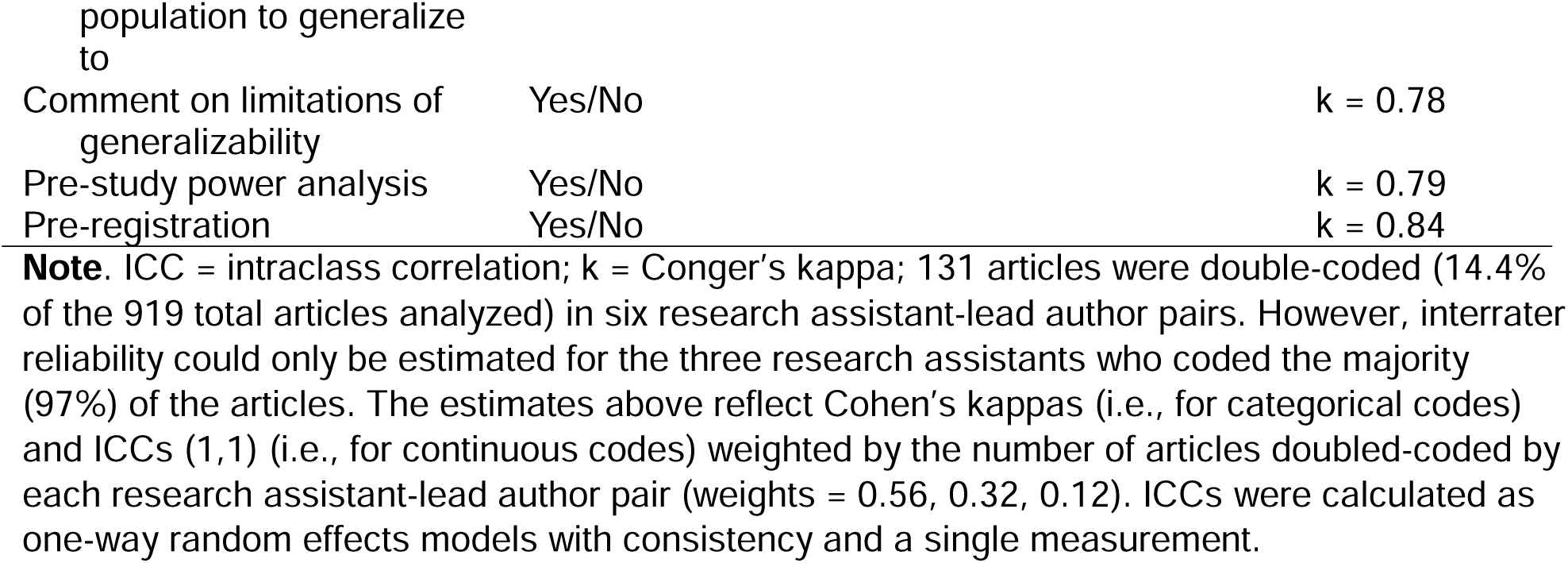
Full-text data extraction codebook and interrater reliability.

To ensure that all data was collected from each article, research assistants coded whether an article cited previously published work to describe the methods. Subsequently, two coders referenced the cited papers to determine if the information was available. If so, [the information in the cited article was used to code that article’s reporting practices]. For demographic information and information about data loss due to poor data quality, the cited paper had to report the exact sample size of the article being coded. Recruitment information and data loss due to participant refusal in the neuroimaging component of the study could source from any cited previous paper using the same participant sample. Referring to previously published work occurred in 14.9% of coded articles with regards to demographic information and 28.4% of coded articles with regards to methodological features.

Articles that passed the abstract and title screening phase were randomized to one of six RAs for full-text data extraction. To ensure reliability in the data extraction phase, the lead author extracted data for a random 15% of articles reviewed by each research assistant (Polanin et al., 2019). Interrater reliability between each RA and the lead author was calculated for every extracted data point using Conger’s Kappa (unweighted) for categorical data and intraclass correlations (one-way random effects, consistency, single measurement) for continuous data (Table 1).

### 2.3. Analytic Plan

All the data and analytic code used in the current paper is freely available at https://osf.io/6tpsh/. The first aim was to document current reporting practices in three domains: (1) demographic characteristics, (2) methods, from recruitment to analysis, and (3) generalizability and open science. In addition to item-level frequencies, we constructed a cumulative reporting score for each domain. The demographics reporting score consisted of four items: race or ethnicity, a measure of socioeconomic resources, sample age, and sex or gender. The methods reporting score included seven items: recruitment strategy, initial recruitment efforts, date of data collection, inclusion/exclusion criteria, reasons for no imaging data, quality control exclusions, and missing values analyses. The generalizability and open science reporting score included four items: identification of a target population to which the study would generalize, description of the limitations of study generalizability, pre-registration, and pre-study power analyses. In the construction of the reporting scores, dichotomous items (e.g., was sample sex or gender composition reported) were coded as 1 = Yes, reported or 0 = No, not reported. Categorical items with more than two options were coded in “strict” or “loose” form, to account for variability in field definitions of what constitutes complete reporting. For example, for sample age, the “strict” reporting score assigned a value of “1” to studies that reported *both* age range and average age, while studies that only reported average age or age range were assigned “0.5” and studies that reporting nothing about sample age were assigned “0”; the “loose” reporting score assigned studies with any information about sample age (i.e., average or range) a value of “1” (see Supplementary Table 3). The main text presents the results of features coded in “strict” form; results for “loose” definitions of reporting are in Supplemental Results.

Second, we identified the study characteristics associated with reporting sociodemographic characteristics, methods, and open science and generalizability practices, using multivariate linear regression. Study characteristics that were used to predict each of the three reporting indices included analytic sample size, sample age group, imaging modality, journal, consortia study status, study type, and whether the study was composed of a patient (versus community) sample. Categorical variables were entered as dummy variables, wherein the reference group was always the largest (e.g., for modality, sMRI studies). We also examined whether study characteristics were associated with analytic sample size (i.e., sample size was the dependent variable) using independent samples t-tests and one way analysis of variance tests for categorical variables. Sample size outliers (+/-3SD) were first winsorized, and then sample size was log-transformed.

Lastly, we examined the extent to which reporting practices in each domain were “valued” in the field. By leveraging the Clarivate’s Cited Reference Search tool (Clarivate, 2023), we constructed a measure of “value” defined as the number of times each article was cited. Citation frequency was recorded in March 2022. Multivariate linear regression models were used to predict citation frequency, with reporting scores, study characteristics, and sample size entered as predictors. As with sample size, citation outliers (+/-3SD) were first winsorized, and then citation frequency was log-transformed following the addition of a constant = 1 (i.e., to account for papers with “0” citations).

## 3. Results

### What were the characteristics of studies included for review?

A total of 3,856 articles were identified through PubMed, with no duplicates (Figure 1). During the title and abstract screen phase, 2,847 articles were excluded because the subjects were non-human (n = 694), the study was a review paper, commentary, systematic review, meta-analysis, case study, or methods paper/technical report (n = 1,342), and/or the neuroimaging modality was not MRI or fMRI (n = 1,772). A total of 1,009 articles were submitted to full-text screening and data extraction, of which another 90 articles were excluded. The final sample size for this structured review was 919 articles.

**Figure 1.**
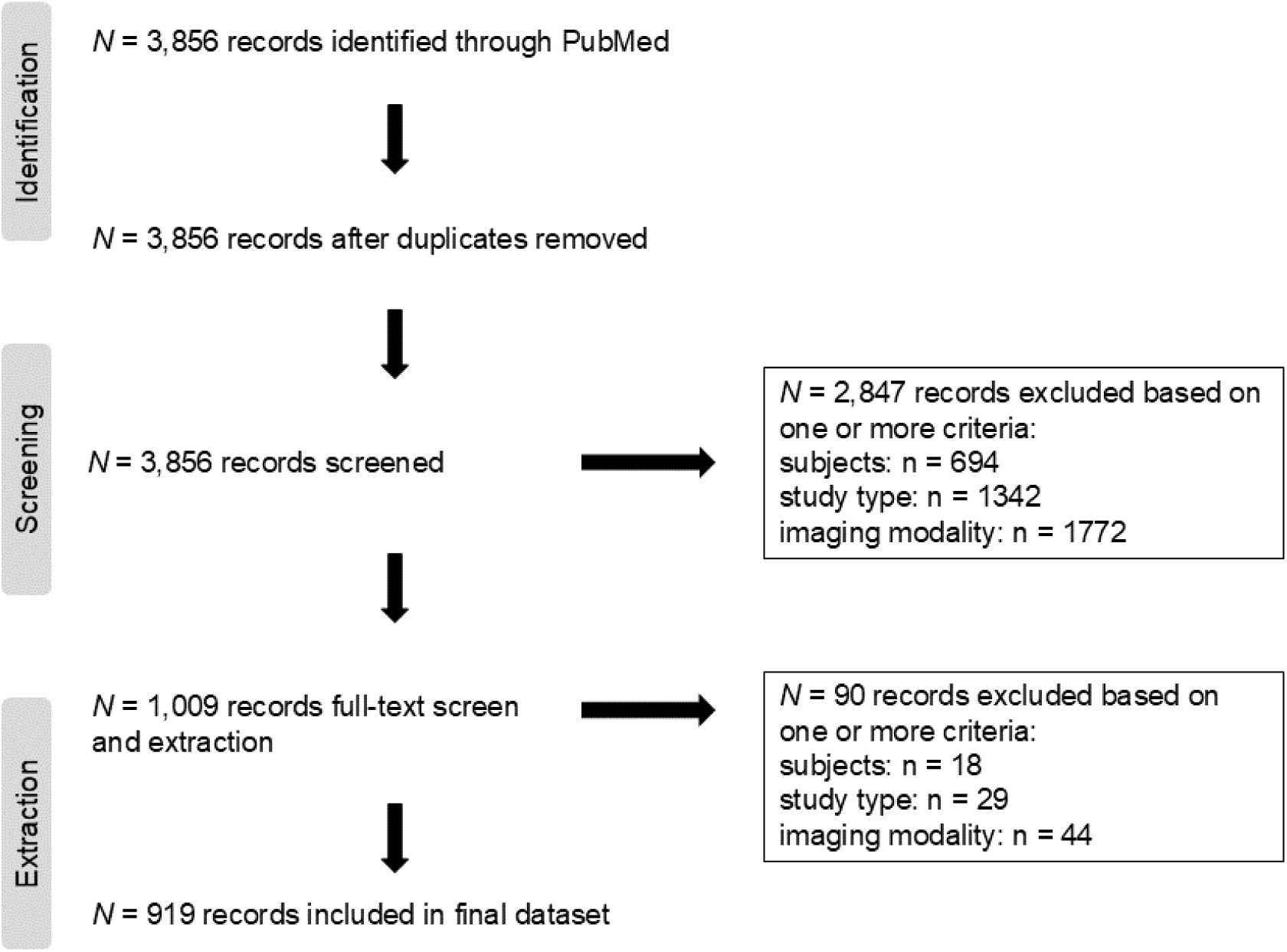
Modified PRISMA Flow Diagram for Study Identification.

Most studies were published in *Neuroimage* (n = 393; 43%) or *Human Brain Mapping* (n = 260; 28%), with the fewest studies published in *Nature Neuroscience* (n = 16; 1.7%) and *Neuron* (n = 7; 0.8%). Nearly 75% of the included studies were conducted in adult samples, followed by child samples (n = 142; 15%) and studies with participants across the lifespan (n = 99; 11%). One-quarter (n = 228) of the studies were classified as patient samples, wherein participants met criteria for a medical diagnosis. Most studies were observational in design (n = 878; 96%) and implemented exclusively fMRI (n = 624; 68%) versus exclusively structural MRI (25%) or combined functional and structural MRI (7.5%). Lastly, 13% of the studies leveraged consortium-level data in that data from multiple sites or studies were combined into a single analysis. See also Supplemental Table 4.

### How large were the included studies and what predicted study sample size?

Across all studies, the largest analytic sample size reported ranged from *N*=5 to *N*=45,615, with an average of *N* = 55 (median) and *N* = 253 (mean). Sample size was associated with other study features such that larger studies were more likely to leverage consortium data (*t*[121.47] = 5.08, *p* < 0.001) and adopt an observational design (*t*[449.85] = 6.35, *p* < 0.001). Sample size was also associated with participant developmental stage (F[2,916] = 17.60, *p* < 0.001) and imaging modality (F[2,916] = 32.34, *p* < 0.001). Post-hoc Tukey tests revealed that studies of participants of all ages were more likely to be larger than studies of children-only (< 18 years) or adult-only participants (mean differences 290.08 and 339.16, respectively; *p*s < 0.001). Functional MRI studies were likely to be smaller in size than both sMRI-only studies (mean difference = 300.45, *p* < 0.001) and multimodal fMRI/sMRI studies (mean difference = 287.22, *p* < 0.001). Analytic sample size was not significantly associated with whether the study examined a patient population (*p* = 0.97). Results were indistinguishable when using the smallest analytic sample size reported (e.g., in cases when the paper reported a sensitivity analysis with a smaller sample size; Supplemental Results).

### Do studies report sociodemographic information about their samples?

Figure 2 displays the proportion of studies that reported demographic, methodological, and generalizability and open science features. Of the studies reviewed (N = 919), 14.8% reported the racial or ethnic identity of participants and 27.9% reported something about the socioeconomic background of participants (e.g., income, education, poverty ratio, Hollingshead score), while 96% reported gender or sex and 98% reported complete (41.7%; both age range and mean age) or partial (56.3%; age range <u>or</u> mean age) information about participant age. Across all four demographic features coded, the average number of features reported per study was two (50% of the total number of features).

**Figure 2.**
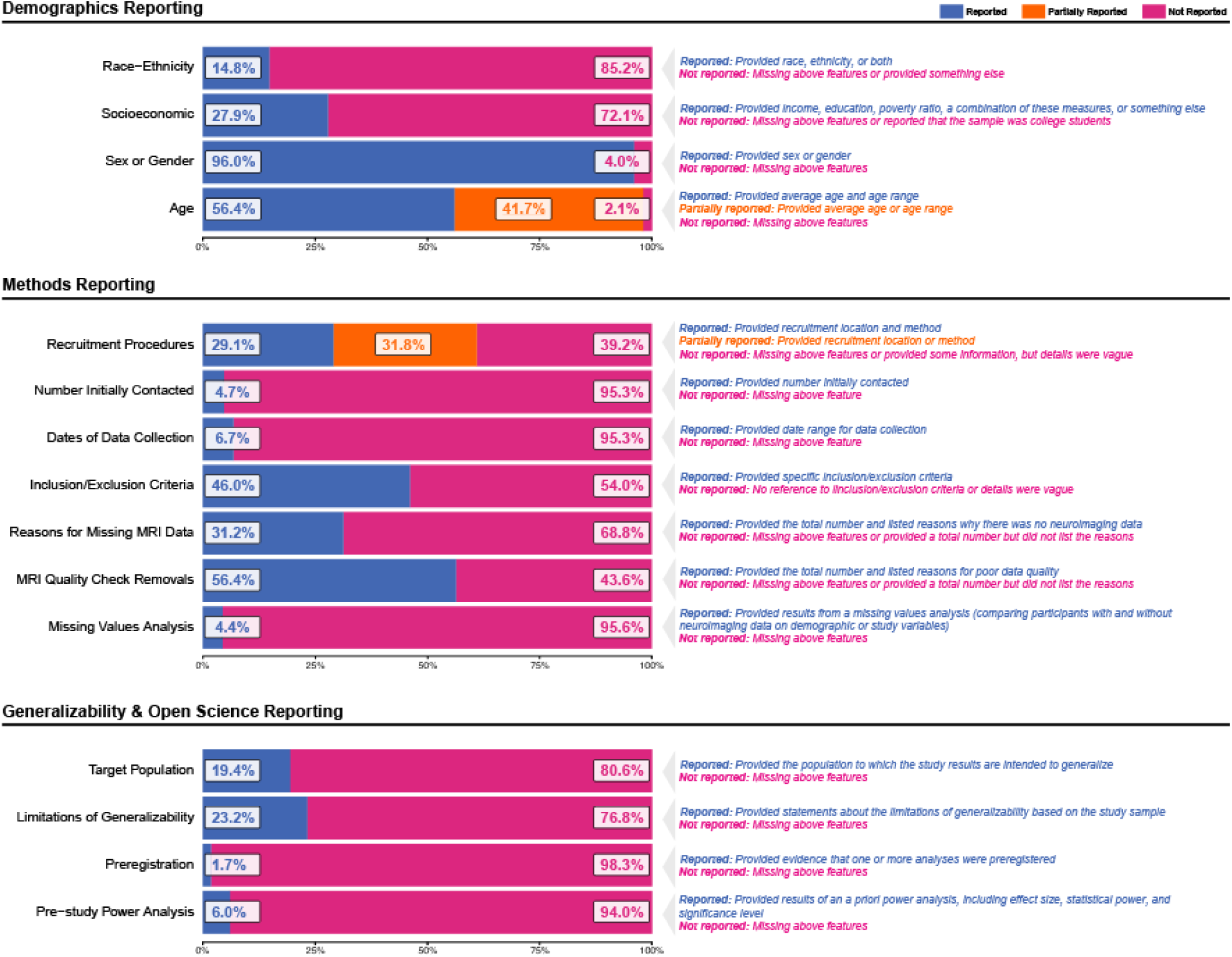
Demographics, Methods, and Generalizability/Open Science Reporting in 919 studies in neuroscience and related journals in 2019. **Note.** *N* = 919. Definitions of whether a feature was reported, partially reported, or not reported can be found in Supplementary Table 2. The proportions shown here reflect “strict” definitions (e.g., for “Age”, complete reporting was operationalized as reporting both age range and mean age, while partial reporting was operationalized as reporting either age range or mean age). See Supplementary Figure 1 for a display of reporting proportions using “loose” definitions.

Within the 136 (∼15%) studies that reported participant racial-ethnic identity in accordance with our codebook, most participants were racialized as White (mean = 56.15%, median = 64.5%), followed by Hispanic/Latinx (mean = 15.10%, median = 6.0%), Asian (mean = 16.65%, median = 0.0%), Black (mean = 12.89%, median = 4.0%), and Biracial or Multiracial (mean = 2.92%, median = 0.0%), with very small proportions of participants racialized as American Indian and Alaskan Native or Hawaiian and Pacific Islander (Figure 3). We acknowledge that these categories are fluid constructs that have changed throughout the historical record and reflect imperfect and incomplete measures of identity and social position (Atkin et al., 2022; Cardenas-Iniguez & Gonzalez, 2024). These racial-ethnic categories are also highly U.S.-centric but, nevertheless, capture the degree to which sociodemographic diversity is represented in the neuroimaging papers reviewed.

Of the 330 (35%) papers that reported something about the socioeconomic background of participants, most (n = 155) reported education continuously (i.e., average education in years) or categorically (e.g., % less than high school degree; % college-educated). The average educational attainment of participants in studies that reported education continuously was 14.98 years (Min – Max = 6 – 21 years), suggesting that, on average, most participants had some education beyond high school or secondary school. As there is wide variability in educational attainment globally (Goujon et al., 2016), we calculated the difference between a study sample’s average education in years and the average years of schooling in the country of recruitment (Figure 4), for the N = 140 studies that reported education continuously and reported the country of recruitment. Eleven studies reported that the study sample was less educated than the average education (in years) in the country of recruitment and 29 studies reported sample education levels within one year of the country average. The remaining 100 studies included participants whose average years of schooling exceeded the country-specific average education level (Figure 4).

### Do studies report the flow of participants from recruitment to final analytic sample?

As with sociodemographic reporting, there was wide variability in the reporting of methodical features related to the flow of participants from recruitment to final analytic sample (Figure 2). Few studies reported the number of participants initially contacted for recruitment (4.7%), the dates of data collection (6.7%), or a missing values analysis comparing participants with and without usable data (4.4%). Far more studies reported information about recruitment procedures – either with complete (38.3%; where participants were recruited from and through what method) or partial (22.5%; where participants were recruited from *or* through what method) information. Forty-six percent of studies reported inclusion/exclusion or eligibility/ineligibility criteria, 31.2% reported reasons for missing imaging data (if applicable) and 56.4% reported the quality control criteria that resulted in the additional exclusion of participants from the analytic sample(s). Across the seven methodical features coded, the average number of features reported per study was two (∼30%).

**Figure 3.**
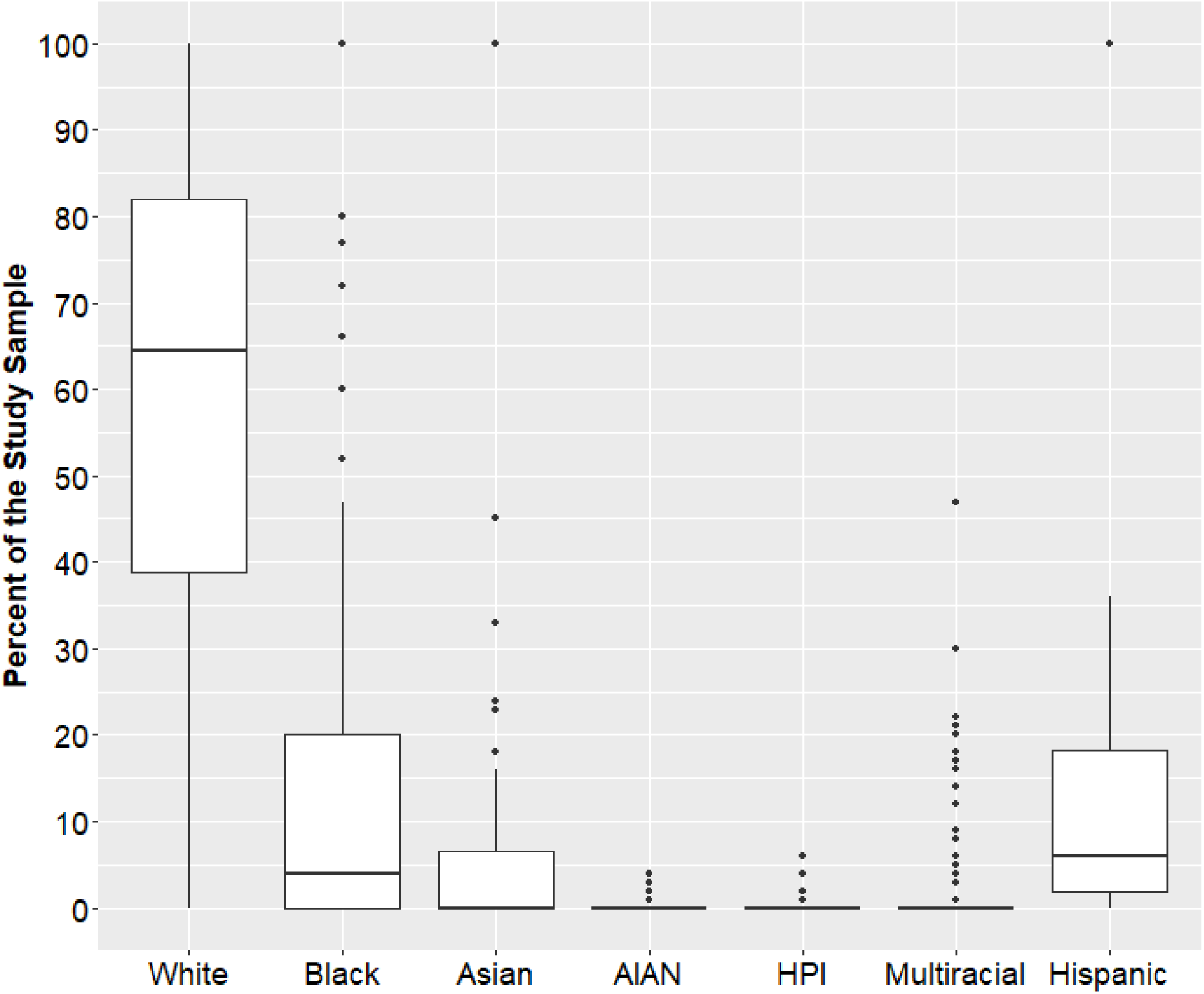
Among studies that report sample race-ethnicity, most participants are racialized as White. **Note**. Includes *N* = 136 (14.8%) studies that reported participant race, ethnicity, or both. Box-plots depict study-specific proportions of participants across seven US-centric racial-ethnic groupings. Note that these figures do not add up to 100% because every study did not report racial-ethnic breakdown for all of the identity categories coded in this structured review.

### Do studies report open science practices?

Lastly, within the open science and generalizability domain, the average number of features reported by each study was zero. Only 19.4% of studies explicitly defined the target population for inference, 23.2% of studies commented on the limitations of the generalizability of their study sample, and 1.7% and 6% of studies pre-registered their aims and hypotheses or conducted a pre-study power analysis, respectively (Figure 2). Results using the alternative “loose” coding scheme were similar (see Supplemental Results).

### Which study characteristics are associated with reporting likelihood?

We next sought to explore the characteristics of studies that report demographic, methodological, and generalizability and open science features. Zero-order correlations indicated that reporting in one domain was associated with reporting in another domain (0.19 < *r* < 0.28, all FDR-adjusted *p*s < 0.001). Sample size was not associated with the demographic, methods, or overall reporting indices, but larger studies were more likely to report study features related to open science and generalizability (*r* = 0.12, FDR-adjusted *p* < 0.001). The same patterns of association were observed using the looser definitions of reporting and the smallest sample size that was reported (Supplemental Results). Next, four multivariate models were estimated, one for each reporting index (demographic, methods, open science and generalizability), as well as the overall reporting index, as the dependent variables (Table 2). All study characteristics and sample size were entered as independent variables. Sample age group, imaging modality, and journal were each uniquely associated with overall reporting frequency. Child and lifespan studies were more likely to report study features than adult-only studies (Overall Reporting Index: Figure 4A), sMRI and multimodal (sMRI/fMRI) studies were more likely to report than fMRI-only studies (Overall Reporting Index: Figure 4B), and studies published in *Developmental Cognitive Neuroscience*, the *American Journal of Psychiatry*, and *Molecular Psychiatry* were more likely to report study features than studies published in *Neuron*, *Journal of Cognitive Neuroscience*, and *Neuroimage* (Overall Reporting Index: Figure 4C).

**Table 2.** Study Characteristics are Associated with Reporting Frequency.

|  | Demographics | Methods | Open Science & Generalizability | Overall Reporting Index |
| --- | --- | --- | --- | --- |
| | B (SE), $\beta$ | B (SE), $\beta$ | B (SE), $\beta$ | B (SE), $\beta$ |
| **Sample Size** | 0.000002<br>(0.00005), 0.001 | -0.00006<br>(0.00008), -0.02 | 0.00007<br>(0.00005), 0.05 | 0.000013<br>(0.00013), 0.003 |
| **Patient Population** | 0.10 (0.06), 0.05 | 0.30 (0.10), 0.11** | 0.16 (0.06), 0.10** | 0.06(0.02), 0.12*** |
| **Consortia Data** | -0.02 (0.08), -0.11** | -0.20 (0.13), -0.05 | 0.10 (0.07), 0.05 | -0.03 (0.02), -0.06 |
| **Treatment / Intervention** | -0.01 (0.01), -0.03 | 0.60 (0.20), 0.10** | 0.09 (0.11), 0.03 | 0.057 (0.03), 0.06 |
| **Sample Age** | F(2, 13.42) =<br>12.10*** | F(2, 40.57) =<br>14.11*** | F(2, 4.61) =<br>4.97** | F(2,147.92) =<br>22.09*** |
| Children | (ref) | (ref) | (ref) | (ref) |
| Adults | -0.40 (0.08),<br>-0.21*** | -0.65 (0.12),<br>-0.22*** | -0.21 (0.07),<br>-0.13** | -1.22 (0.02),<br>-0.27*** |
| All ages | -0.20 (0.10), -0.06 | -0.30 (0.16), -0.07 | -0.06 (0.09), -0.03 | -0.05 (0.02), -0.08* |
| **Modality** | F(2,19.83) =<br>17.87*** | F(2,1.28) =<br>0.45 | F(2,5.64) =<br>6.08** | F(2,62.75) =<br>9.37*** |
| Structural MRI | (ref) | (ref) | (ref) | (ref) |
| Functional MRI | -0.40 (0.06),<br>-0.22*** | -0.09 (0.10),<br>-0.03 | -0.20 (0.06),<br>-0.13*** | -0.067 (0.02),<br>-0.16*** |
| Multimodal | -0.30 (0.01),<br>-0.09* | -0.020 (0.17),<br>-0.005 | -0.10 (0.10),<br>-0.04 | -0.038 (0.03),<br>-0.05 |
| **Journal** | F(8,5.58) =<br>1.26 | F(8,40.61) =<br>3.53*** | F(2,4.31) =<br>1.16 | F(8,70.92) =<br>2.65** |
| *NeuroImage* | (ref) | (ref) | (ref) | (ref) |
| *DCN* | -0.01 (0.01), -0.004 | 0.55 (0.18), 0.11** | -0.05 (0.10), -0.02 | 0.049 (0.03), 0.06 |
| *HBM* | -0.08 (0.06), 0.04 | 0.23 (0.10), 0.08* | -0.06 (0.06), -0.04 | 0.025 (0.02), 0.06 |
| *JOCN* | -0.09 (0.01), -0.03 | -0.10 (0.17), 0.02 | -0.08 (0.10), -0.03 | -0.01 (0.03), -0.01 |
| *Mol Psych* | 0.10 (0.01), 0.02 | 0.47 (0.23), 0.07* | 0.17 (0.13), 0.04 | 0.07 (0.04), 0.07* |
| *Nature Neuro* | -0.30 (0.20), -0.04 | 0.96 (0.31), 0.10** | 0.15 (0.18), 0.003 | 0.07 (0.05), 0.05 |
| *Neuron* | -0.30 (0.03), -0.03 | -0.29 (0.46), -0.02 | -0.35 (0.26), -0.04 | -0.09 (0.07), -0.04 |
| *SCAN* | 0.15 (0.09), 0.05 | 0.41 (0.15), 0.09** | 0.13 (0.09), 0.05 | 0.07 (0.02), 0.10** |
| *AJP* | 0.24 (0.02), 0.05 | 0.72 (0.29), 0.08* | 0.05 (0.16), 0.009 | 1.01 (0.04), 0.07* |

*Note.* $N = 910$ ; Sample size refers to the largest sample size reported, which was first winsorized to +3SD from the mean, and log-transformed. Multimodal refers to studies that included both structural and functional MRI. One-way ANOVAs for categorical variables evaluate the change in model fit when that predictor was removed from the model, compared to the full model. The reporting indices here reflect “strict” definitions (see Methods); Supplemental

**Figure 4.**
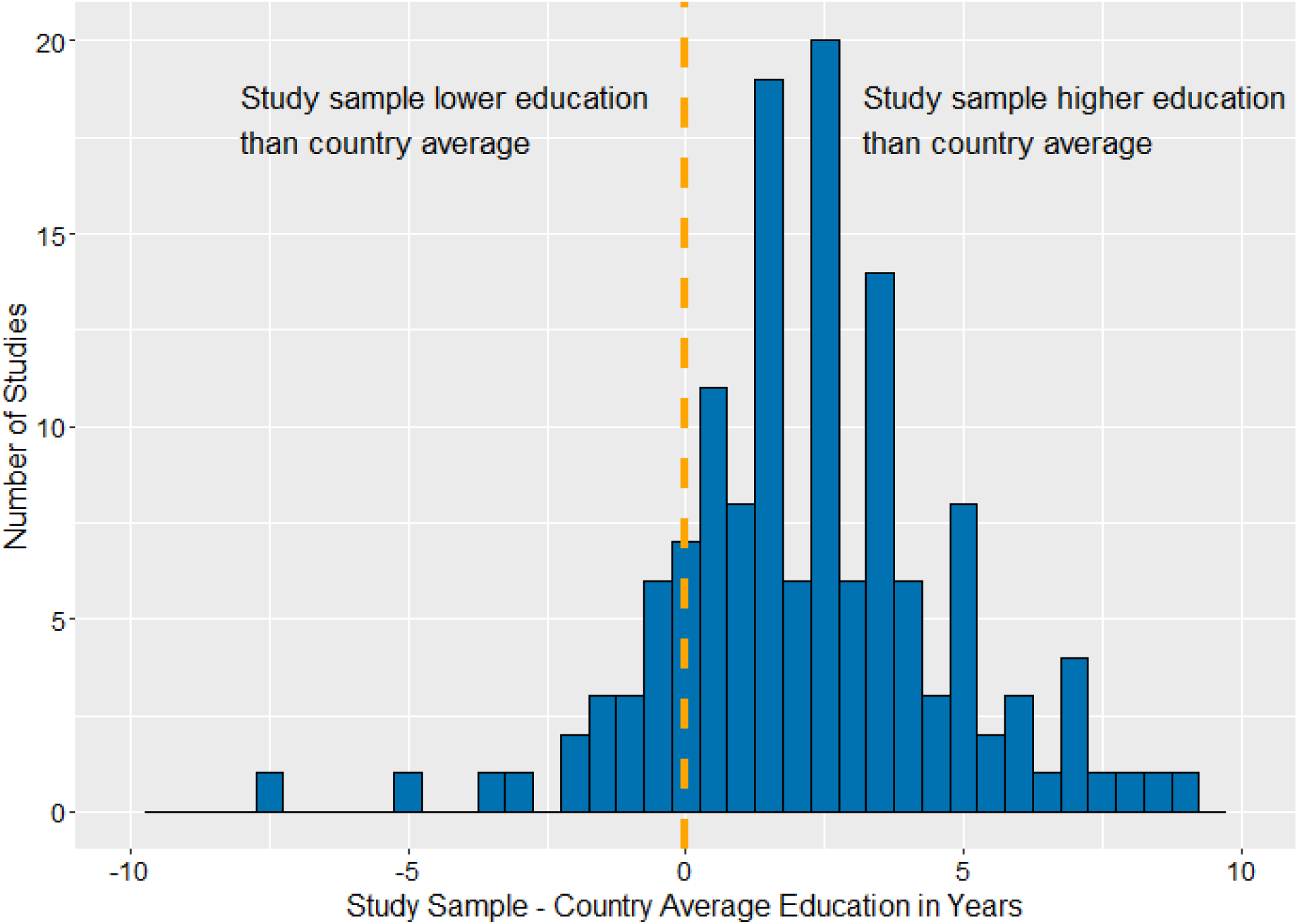
Studies overrepresent highly educated participants relative to the recruitment country average years of schooling. **Note**. Of the 140 (15.2%) studies that reported both country of recruitment (n = 317 [34.5%]) and mean education in years (n = 155 [16.9%]), average education of the study samples recruited in each country is plotted against that country’s average education in years. Country population estimates were drawn from Our World in Data (https://ourworldindata.org/global-education). Studies that included several countries of recruitment were excluded from this analysis (N = 7). Nineteen countries are represented in this figure.

**Figure 5.**
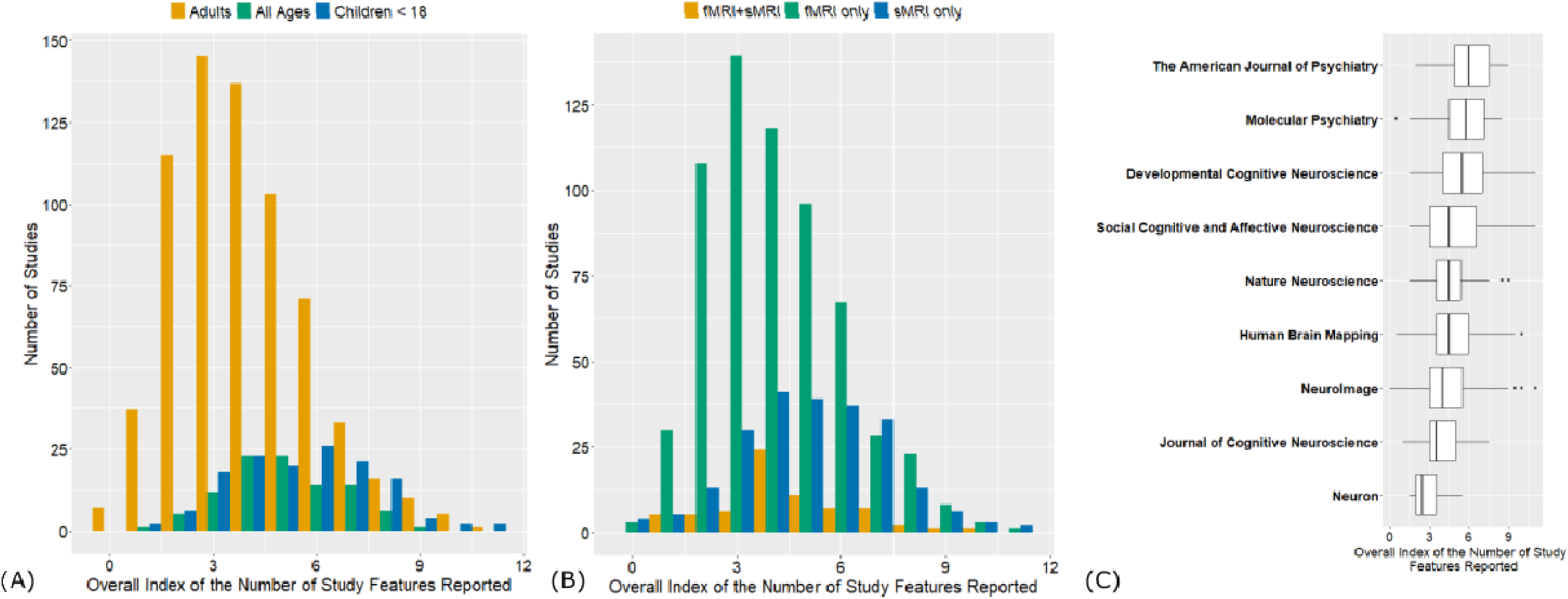
Sample age, imaging modality, and journal are associated with overall reporting frequency. **Note**. *N* = 919. (A) Overall index of the number of study features reported (out of a possible 15 features), plotted by the reported sample age group. Adult-only studies were less likely than both child-only and lifespan studies to report study features across domains (Table 2). (B) Overall index of the number of study features reported, plotted by imaging modality. Studies employing multimodal or structural MRI only were more likely to report studies features across domains than functional MRI only studies (Table 2). (C) Overall index of the number of study features reported, plotted by journal (see Table 2 for multivariate results)

Patterns were similar using the domain-specific (sociodemographic, methods, or open science and generalizability) reporting indices (Table 2). Of note, non-consortia studies were more likely than consortia studies to report demographic features, and studies using patient samples were more likely to report methods and open science and generalizability features than studies that did not recruit patients. Consistent with the zero-order associations, sample size was not associated with the overall, sociodemographic, or methods reporting indices, and was no longer associated with reporting in the open science and generalizability domain after study characteristics were included in the multivariate models. Multivariate results were comparable in terms of both strength and direction using the loose reporting criteria (Supplemental Table 4).

### What are the associations between reporting frequency, sample size, and citation counts?

Given the ubiquity of reporting standards across research fields (e.g., APA, STROBE, EQUATOR) combined with the wide variability in reporting identified here, the last empirical exercise evaluated the extent to which reporting features within and across domains were valued in terms of citation count within two years of publication. Operationalizing “value” through citation frequency is an imperfect measure and one with biases including weak prediction of research quality (Dougherty & Horne, 2022). At the same time, more frequently cited papers are also likely more visible to both the research community and the public (McKiernan et al., 2019; Sternberg, 2016). In the ∼two years after publication (January 2020 – March 2022), the number of times each paper was cited ranged from 0 to 112 (Mean [SD] = 11.51[12.15], Median = 8).

After accounting for all study characteristics (i.e., journal, participant age group, modality, study type, consortia status, patient population) and sample size, none of the domain-specific reporting scores (demographics index: B[SE] = −0.004[0.032], *p* = 0.90; methods index: B[SE] = −0.016[0.021], *p* = 0.43; open science and generalizability index: B[SE] = 0.014[0.035], *p* = 0.68) were associated with citation frequency. By contrast, study sample size was strongly associated with citation frequency (B[SE] = 0.127[0.025], *p* < 0.001)^1^. For every one unit increase in sample size, citation frequency increased by 12.7%. This figure was consistent in models with the overall reporting index, as well as using the loosely defined transparency scores (see Supplemental Results).

## 4. Discussion

In the last 30 years, advances in MRI and fMRI technology have enabled scientists to study the brain bases of human behavior, cognition, and health and disease. In turn, human neuroscience has captured the attention of scientists, policymakers, and the public at large. Metascience inquiries are needed to evaluate the state of this field, with particular emphasis on the generalizability and reproducibility of study findings. And yet, to evaluate the rigor of a science, researchers must report study demographics, methodical decisions, and adherence to principles of open science. This structured review of current methodological practices in human MRI and fMRI studies documented reporting practices across three domains (i.e., demographic, methods, open science and generalizability). Results indicated that although some study features were widely reported (e.g., gender or sex), many were not (e.g., reasons for missing imaging data). This pattern was observed in all three reporting domains, and reporting in one domain was associated with reporting in others. Coded study characteristics such as sample age group, journal, and imaging modality were related to a study’s likelihood of reporting across domains. Moreover, when examining associations between study characteristics and citation frequency, study sample size, but not reporting frequency, was strongly associated with greater visibility in the field.

Two surprising results were low reporting rates of inclusion/exclusion criteria and reasons for missing MRI or fMRI data. Without this information, which is clearly specified in existing reporting guidelines (American Psychological Association, 2020; Elm et al., 2008), it is impossible to know the characteristics of the population that are not represented in a given study. More than half (54%) of included studies in this structured review did not explicitly list inclusion/exclusion or eligibility/ineligibility criteria. In a qualitative review of these studies, we found widespread non-specific statements such as “All participants were healthy and right-handed*”* or “Participants were free of major psychiatric disorders*”.* On the other hand, studies with clear reporting included numbered lists of exclusion criteria and definitions for each criterion. For example, van Harmelen and colleagues (2014) defined “infrequent benzodiazepine use” as “the equivalent to two doses of 10 mg of oxazepam three times per week or use within 48 h prior to scanning” (van Harmelen et al., 2014). An even larger proportion of studies (68.8%) failed to report the contributions of missing MRI or fMRI data to the “flow of participants” (American Psychological Association, 2020). More often, studies stated the number of participants used in the current analysis without reference to any of the reasons why participants did not have imaging data (and did not state whether participation in the MRI scan was an inclusion or eligibility criterion). This is a key piece of missing information that prevents advancements in the field. By identifying characteristics of individuals who do not enter the scanning environment (e.g., anxiety or hesitation related to the scan), researchers can adjust protocols to promote retention (e.g., mock scanners, hiring staff with similar experiences to participants) (Charpentier et al., 2021; Gard et al., 2023; Perlman, 2012). One excellent example of reporting the flow of participants can be found in Hahm et al. (2019), who provided a detailed account of how participant data was lost at every stage of filtering, from the original sample size (N = 3,309) to the final analytic sample size (N = 3,027). Another way to present this information is through the use of a dropout table that includes reasons for no scanning and reasons for analytic exclusion based on data quality decisions (e.g., Gard et al., 2021).

Another concerning finding from the current study was the low reporting of select demographic characteristics, including the racial-ethnic identity and socioeconomic background of participants. Race-ethnicity is not biological construct; it reflects a constellation of environmental, personal, and community experiences, including historical and current structural racism (Cardenas-Iniguez & Gonzalez, 2024; Varnum et al., 2010; Webb & Harnett, 2024; Williams & Collins, 2001). Racial-ethnic identity is an especially salient sociodemographic characteristic that describes participants’ lived experiences and leads to culturally-specific trajectories of development (Iruka et al., 2022). In turn, a cadre of studies have shown that racialized (and socioeconomic) experiences shape brain function and structure (Harnett et al., 2024; Hyde et al., 2020; Monk & Hardi, 2023; Webb & Harnett, 2024), trauma and memory (Roozendaal et al., 2009), socioeconomic resources and executive function (Palacios-Barrios & Hanson, 2019), and caregiving and emotion processing (Farber et al., 2020). Racialized experiences also include promotive factors (e.g., ethnic-racial identity, community cohesion) that have been linked to brain development. For example, in a sample of diverse young adults, greater ethnic identity resolution (i.e., having clarity about what is means to be a member of a racial-ethnic group), which is routinely higher in racial-ethnic minoritized youth and associated with adaptive outcomes (Umaña-Taylor et al., 2014), was associated with higher frontoparietal network density (Constante et al., 2023). Thus, studies that ignore racialized experiences and environmental factors related to the provision of economic resources (e.g., Hyde et al., 2020) may neglect contributors to heterogeneity in brain development. Indeed, a recent investigation using a large study of early adolescent youth reported greater heterogeneity in neurodevelopment within marginalized and structurally disadvantaged groups compared to their more advantaged peers (Bottenhorn et al., 2024). An underexplored hypothesis is that greater heterogeneity accounts for recent findings (Greene et al., 2022) that brain-based predictive models are less prognostic for individuals who defy sample averages (e.g., high cognitive scores despite lower educational opportunity). By understanding heterogeneity within sociodemographic groups, the field can begin to integrate strengths-based neuroscience research questions.

That human neuroscience emerged from animal neuroscience may help to explain why reporting rates of sample sociodemographic features are so low. All humans, regardless of social status, resources, identity, and/or geographical location, share fundamental biological processes. Dendritic arborization, cell death, synaptic pruning, and myelination are examples of brain-based molecular processes that function in the same way across all humans. However, in contrast to animal neuroscience, human neuroimaging studies do not measure processes at a cellular level. Rather, our indirect measures of brain structure and function capture multiple molecular processes simultaneously. And as our level of inference shifted from molecules and cells in animal neuroscience to individuals and groups of people in human neuroscience, so too did the importance of attending to variation in environmental contexts (Falk et al., 2013). In short, one brain is not representative of all brains. However, among the studies included in this structured review, most (76.8%) did not comment on the limitations of sample generalizability or identify the target population to which the study sought to generalize to (80.6%). Instead, many studies invoked generic language (DeJesus et al., 2019) that implies universalisms (e.g., “Furthermore, these results suggest that the right IFG plays a crucial role in supporting pitch encoding in the typical brain”).

A related result from this review is the recognition of the large role that study sample size plays in both the interpretations of generalizability and in the visibility of research products. Qualitatively, there were several examples of falsely equating sample size with population representation. For example, one discussion section read, “Since our sample size was comparatively larger than in previous studies and included 2,400 measurements for each session, the found partial correlations can be seen as more representative of the true effect sizes in the population”. Larger sample sizes indeed enable greater statistical power to detect small effect sizes, thus contributing to scientific replicability (Bossier et al., 2020; Marek et al., 2022; Turner et al., 2018). However, the notion that sample size in and of itself is a marker of generalizability is faulty at best. Decades of research in survey methodology has highlighted the insidious impact of non-response bias on deriving population estimates (Groves et al., 2009; Heeringa et al., 2017). In one applied example, Bradley and colleagues (2021) compared millions of survey responses from three COVID-19 symptom trackers to benchmark vaccine uptake statistics from the Centers for Disease Control and Prevention in the United States. Results indicated that survey studies significant overestimated vaccine uptake by 11 to 20%, despite boasting sample sizes large enough to detect the true effect (Bradley et al., 2021). The lack of population generalizability in large sample sizes has also been demonstrated with neuroimaging data (Gard et al., 2023; LeWinn et al., 2017). For example, LeWinn and colleagues (2017) adjusted the sample composition of a large community-based neuroimaging sample (N = 1,162) to match the demographics of the target population of children in the United States; this process increased the contribution of Hispanic children and children of caregivers with a high school education or less to the overall test statistics. The unweighted sample produced an s-shaped curve of age-related associations with total cortical volume, while the weighted (and more representative) sample produced a quadratic association between age and total cortical volume (LeWinn et al., 2017). Were these results to be used as benchmark values, in the same way as height charts are used in pediatrician offices, some children would be misclassified as on or off developmental targets. Larger sample sizes may not produce inferences that generalize to a broader population of individuals. And yet, results from the current study also made clear that reporting larger sample sizes was highly “valued” by the field (i.e., in terms of citation count), as compared to reporting study sample demographics, methodological decisions, and adherence to principles of open science and generalizability. Estimates from multivariate models that accounted for coded study features (e.g., journal, imaging modality) demonstrated that for every additional 1-unit increase in study sample size, paper citations increased by nearly 13%. By contrast, the association between reporting frequency and citation count was estimated at a precise zero.

### Recommendations

To more critically evaluate the state of the field and to generate scientific advances in technology and participant retention strategies, we must increase reporting transparency in human neuroscience studies. In this review, studies that featured adult-aged participants, implemented fMRI, and/or leveraged consortia-level data were less likely to report across domains. Although individual studies and researchers could be targeted for intervention, intervention at the journal-level might prove more effective and efficient for improving reporting culture. Indeed, such structural requirements already exist; for example, journals within the *Nature* publishing group require authors to complete an editorial policy checklist that includes a data availability statement and confirmation of practices for the protection of research participants and biological samples. Forms like these, which authors can complete upon manuscript submission to a journal, could be expanded to include information about sample demographics, methodological details, and open science and generalizability practices.

In turn, we propose a journal-level intervention to bolster reporting practices in human neuroimaging studies through a reporting checklist that authors complete at the time of manuscript submission (Table 3). Checklist items are divided into the three domains of reporting that we evaluated in this review – sample sociodemographic characteristics, methodological features, and open science and generalizability practices. For each item, authors select whether the manuscript includes the information, just as current submission checklists require authors to acknowledge that all authors have approved the manuscript. Importantly, even if journals do not have the capacity to enforce all reporting features, the simple act of acknowledging whether said information is included in the manuscript may encourage researchers to increase their reporting practices more widely. In some cases, though not ideal, researchers can also acknowledge that the requested information was not collected (e.g., reasons for ineligibility).

**Table 3.** Proposed submission reporting checklist to facilitate increased methodological transparency in human neuroimaging research.

| **Sociodemographic Features** | **Additional Guidance** |
| --- | --- |
| Participant age | Report average, standard deviation, and range |
| Gender identity and/or sex | Gender identity (current); Biological sex (male/female/intersex) at birth. Both offer distinct pieces of information. |
| Community of descent | Context-specific. Any measure that captures some aspect of heritage or identity. Examples using race-ethnicity, ethnicity, country of origin, language. These are not intended to be categories based in biology but, rather, as markers of identities imposed in each context that conveys something about representation. |
| Socioeconomic resources | Any measure that captures social and/or economic resources. Examples include income, education in years, area-level measures of resources |
| **Methodological** |  |
| Recruitment procedures | Include both the location (e.g., schools) and method (e.g., flyers to all students). In studies of cases/controls, report whether recruitment procedures differed |
| Dates of data collection |  |
| Inclusion and exclusion criteria | Use the standard terminology of inclusion/exclusion or ineligible/eligible for clarity |
| Flow of participants from recruitment through analytic sample | Number of participants who completed each of the following: (1) initial recruitment, (2) completed imaging protocol, (3) had high quality imaging data, (4) had acceptable data according to other non-imaging analytic decisions |
| Missing data analysis | Comparison of the analytic sample and original sample on sociodemographic constructs and primary variables of interest. |
| Reasons for data loss | List specific criteria (e.g., high motion; > 1mm framewise displacement) to identify sources of data loss due to incomplete imaging data and low-quality imaging data |
| **Open Science & Generalizability** |  |
| State intended target population | What are the characteristics of the target population to which the study intends to generalize? |
| Comment of limitations of generalizability | Are there features of the study design and the recruited participants that temper population generalizability? |
| Were any pre-study power analyses conducted? | If applicable, clarify that pre-study power analyses were conducted prior to study implementation |
| Was any part of the analysis pre-registered | If applicable, state and provide link |
| Data and/or code availability | If applicable, state and provide link |

One major challenge to this proposal is that racial-ethnic identity is arguably a difficult study feature to code and compare worldwide. For example, in France it is illegal to collect demographic information about an individual’s race or ethnicity (Léonard, 2014). Here, we endorse a recommendation advanced by the ManyBabies international consortium for studies to adopt a measure of community of descent that is locally valid and capture aspects of heritage and identity (Singh et al., 2023). Multiple constructs are captured under the term “community of descent” proposed by Singh and colleagues (2023), including ancestry, race-ethnicity, religion, national origin, cultural practices, and native language use, among others.

Reporting sample sociodemographic characteristics must go hand in hand with reporting methodological decisions (e.g., inclusion/exclusion criteria, data quality restrictions) and adherence to principles of open science and generalizability (e.g., preregistration of exploratory or confirmatory hypotheses, pre-study power analyses, identification of a target population). The composition of a study sample is related to how participants were recruited and retained, and how the data was cleaned and analyzed. Several comprehensive reviews highlight the multitude of tools available to researchers for promoting the reproducibility and replicability of human neuroscience research (e.g., Klapwijk et al., 2021). Results from this structured review suggest that journal-level interventions can augment adjustments made by individual researchers.

### Limitations

Despite the strengths of this structured review, several limitations warrant consideration. First, this review cannot address current methodological practices in neuroimaging modalities beyond MRI or fMRI. Second, articles whose primary purpose was to evaluate a new method or technique, rather than generalize results to a population, were also excluded. Although one may argue that methods papers should also be held to reporting standards, we excluded these papers so as not to bias our results; an initial review of methods papers and technical reports revealed that few reported detailed methods.

### Future Directions and Conclusions

This metascience investigation of reporting practices in human neuroscience studies contributes to our understanding of scientific rigor in the field, including generalizability and reproducibility. However, the question of whether and how to report methodological practices is distinct from efforts to increase diversity in human neuroimaging research. The former can be implemented now (Table 3); the latter requires specialized training, experience, and resources (Habibi et al., 2015; Roosa et al., 2008). Supporting researchers with specialized skills to recruit and retain communities historically excluded from scientific research is essential to increasing the generalizability of our science. Ultimately, efforts to increase sample sociodemographic representation will require radical shifts in how we conduct neuroimaging studies; such a shift requires time and patience to rebuild relationships with communities historically wronged by scientific studies (Foster et al., 2024; Gard et al., 2022; La Scala et al., 2023).

The “Age of the Brain” is very much upon us; human neuroscience informs public life in policy, medicine, and public health domains. At the same time, the opportunity to use human neuroimaging data to understand human behavior and support human wellbeing comes with responsibility to participants, funders, and the public at-large. Promoting greater transparency in reporting practices is one of many efforts that are needed to gain a more generalizable, reliable, and reproducible understanding of the human brain across environmental contexts.

## Footnotes

1 As both citation frequency and study sample size were log-transformed variables, unstandardized estimates can be interpreted as percent change.

## Notes

https://osf.io/6tpsh/

### Competing Interest Statement

The authors have declared no competing interest.

