## Supplemental Material for "Current Reporting Practices in Human Neuroscience Research"

**SUPPLEMENTARY MATERIAL**

**Supplemental Methods**

**Coder Training Procedures**

During orientation, undergraduate and postbaccalaureate research assistants (RAs) were trained in best practices for conducting systematic reviews (Polanin et al., 2019), project goals, the coding manual, and how to use the systematic review software DistillerSR (*DistillerSR*, 2023). Next, 25 randomly selected neuroimaging articles published in 2018 from target journals were selected for training. All RAs and the lead author screened the 25 articles according to the exclusion criteria set forth in the coding manual. Discrepancies were discussed as a group, and clarifications were added to the coding manual. Next, RAs and the lead author extracted data from 10 articles according to the coding manual. Discrepancies were discussed as a group. Additional edits to the coding manual were added to improve clarification and reliability. RAs then independently coded at least 5 additional articles. RAs were “approved” for independent coding once they reached 90% agreement with the lead author. After coding approval, RAs were instructed to work in blocks of 2 hours of less; research suggests that reliability wanes when reviewers complete screening and data extraction across blocks of time longer than 2 – 3 hours (Polanin et al., 2019). Three RAs plus the lead author conducted the title and abstract screening (N = 3856 articles), and three RAs (one newly added, as another RA graduated) conducted the stage two full-text data extraction (N = 1010 articles).

**Deviations from Preregistered Protocol**

Although the coding manual and procedures were preregistered prior to data collection and analysis ([https://osf.io/6tpsh/](about:blank)), the training and initial screening phases of the project revealed several concerns for the viability of completing the structured review in a timely manner. Following preregistration, we decided to exclude articles that implemented any neuroimaging modality except for MRI and fMRI (i.e., fNIRS, EEG, MEG, SPECT, TMS). Second, we simplified the data extraction form by removing several questions and adding additional questions to streamline the full-text data extraction (Supplemental Table 1). In addition, although the preregistered coding manual stated RAs would screen the title and abstracts of all articles followed by 15% reliability check by the lead author, two RAs reviewed every title and abstract for inclusion, and discrepancies were adjudicated by the first author. This approach was important for engaging the RAs in the overall goals of the project and served to build rapport within the coding team. Full-text data extraction proceeded as preregistered, with articles randomized to RAs and 15% of articles were checked for reliability by the first author.

**Supplemental Results**

**Alternative Coding Scheme for Reporting Frequency**

In the construction of the reporting scores, categorical items with more than two options were coded in “strict” or “loose” form, to account for variability in field definitions of what constitutes complete reporting (Supplemental Table 3). The codes that were impacted by this process were: (1) *Does the study report the race and/or ethnicity of the sample?* (2) *Does the study report the mean age (in years or months) and the age range of the sample?* (3) *Does the study report the socioeconomic background of the sample?* (4) *To what extent does the study report recruitment procedures?* (5) *Does the study report clear inclusion and exclusion criteria for participation in the study?* (6) *Does the study report the number of participants (if any) refused or were unable to complete the neuroimaging protocol?* and (7) *Does the study report the number of participants (if any) were removed from the analysis due to poor data quality?* Supplemental Figure 1 depicts the results of the looser reporting definitions for these and all codes. In addition to studies that reported participant race and/or ethnicity (14.8%; Figure 3), an additional 24 studies (2.6%) reported “something else” about the participant’s racial-ethnic or cultural identity. Examples included the sample percentage of “Dutch ethnicity” (refid = 31299479), the sample percentage born outside of the United States (refid = 31600019), and genetic ancestry (refid = 31441562). In the loose reporting score for demographics, studies that reporting “something else” about the participant’s racial-ethnic background were considered complete reporting. Also in the loose reporting score for demographics, no distinction was made between studies that reported participant age as range, with mean and standard deviation, or both (97.9%). Lastly, a looser definition of reporting for socioeconomic background included studies that explicitly recruited college samples (n = 33; 3.6%). Although we do not yet know whether participants in college will complete their schooling, recruiting college students means that the researchers are implicitly collecting information about participant’s socioeconomic resources. Thus, the overall proportion of studies reporting the socioeconomic resources of the participants changed from 27.9% to 35%. Still, consistent with the strict demographics reporting index, studies reported an average of 2 or 50% of the demographics reporting criteria. In terms of methodological reporting, 60 (6.5%) additional studies with multiple recruitment sites were classified as “complete” reporting, even though information about only one site was given. We defined explicit mention of inclusion/exclusion criteria if the words “inclusion” “exclusion” “eligible” or “ineligible” were used in the written text. There were many more instances of generic language that prevented readers from easily identifying the inclusion/exclusion criteria; the looser coding scheme counted generic statements as partial reporting. Lastly, the looser coding scheme classified studies as partial reporting in cases where a total number, but no reasons, were provided for the two items regarding removing participants because no scan was completed and removing participants due to poor data quality (Supplemental Table 2). The alternative coding scheme resulted in an additional 31.2% and 56.4% of studies that reported information about missing data loss due to no MRI and missing data loss due to poor data quality, respectively (Supplemental Figure 1).

**Characteristics of Reporting Using an Alternative Coding Scheme**

Using the alternative coding scheme with looser definitions of “complete reporting” (Supplemental Table 2), zero-order correlations indicated that reporting in one domain was associated with reporting in another domain (.18 < *r* < 0.27, all FDR-adjusted *p*s < 0.001). Sample size (the largest sample size reported) was not associated with the demographic, methods, or overall reporting indices, but larger studies were more likely to report study features related to open science and generalizability (*r* = 0.12, FDR-adjusted *p* < 0.001).

**Smallest Sample Size**

**Associations between Smallest Sample Size and Study Characteristics.** In addition to coding the largest sample size reported in each study, we also coded the smallest sample size reported in each study. Multiple sample sizes were often reported when authors conducted sensitivity analyses using a subsample of the larger study. Across all studies, the smallest analytic sample size reported ranged from *N*=5 to *N*=5,967, with an average of *N* = 145 (mean) and *N* = 43 (median). Sample size was associated with other study features such that larger studies were more likely to leverage consortium data (*t*[125.20] = 7.39, *p* < 0.001) and adopt an observational design (*t*[117.99] = 6.15, *p* < 0.001). Sample size was also associated with participant developmental stage (F[2,916] = 13.92, *p* < 0.001) and imaging modality (F[2,916] = 33.34, *p* < 0.001). Post-hoc Tukey tests revealed that studies of participants of all ages were more likely to be larger than studies of children-only (mean difference = 83.27, *p* = 0.02) or adult-only participants (mean difference = 125.11, *p* < 0.001). Functional MRI studies were likely to be smaller in size than both sMRI-only studies (mean difference = 128.08, *p* < 0.001) and multimodal fMRI/sMRI studies (mean difference = 128.78, *p* < 0.001). Analytic sample size was not related to whether the study examined a patient population (*p* = 0.96).

**Associations between Smallest Sample Size and Reporting Frequency.** The smallest sample size reported was also associated with overall reporting frequency and reporting in certain domains, regardless of whether “strict” or “loose” definitions were used to construct the reporting indices. In zero-order models with FDR-adjustment of *p*-values, the smallest sample size reported was associated with greater reporting of methodological features (*r* = 0.15, *p* < 0.001) and open science and generalizability features (*r* = 0.15, *p* < 0.001), but not the overall reporting frequency (*r* = 0.05, *p* > 0.10), or methods reporting (*r* = 0.06, *p* > 0.10).

***Supplementary Table 1. Modifications to preregistered coding protocol***

| **Questions that were added to the data extraction form** | **Questions that were removed from or edited within the data extraction form** | **Rationale** |
| --- | --- | --- |
| Should this study be excluded for review? If so, for what reason? |  | To exclude articles that passed the title and abstract screening phase, but were considered ineligible in the full-text data extraction |
| Is this a patient population? [subjects meet diagnostic criteria for a medical disorder] |  | Restrictive recruitment procedures suggest that studies of patient populations should be considered separately. |
| How many participant samples were included in the study? |  | Several papers replicated their results in more than one sample. Only the primary sample was coded |
| Did this study utilize data from a consortium of several participant samples? (e.g., ABIDE, ENIGMA) |  | As data is aggregated across several samples, there is no “primary sample”; thus, consortium papers should be considered separately. |
| Did the study reference a previous paper to report the demographic information? |  | To streamline full-text data extraction, we added this question and returned to papers that references a previous paper. |
| Did the study reference a previous paper to report the recruitment procedures and/or data availability information of the sample? |  | To streamline full-text data extraction, we added this question and returned to papers that references a previous paper. |
|  | Does the study report whether the sample is urban or rural? | This question could not be systematically coded. |
|  | Did participants self-select into the sample/ was the study based on a non-probability sample? | This question could not be systematically coded. |
|  | Does the study report a date range for *neuroimaging* data collection? | To reduce coding burden, we removed this question and retained the question about the date range for the overall data collection |
|  | If the study reported a date range for neuroimaging data collection, what year(s) was the data collected? | This question was removed to streamline full-text data extraction |
|  | Was participation in the neuroimaging component of the study an inclusion criterion? | This question could not be systematically coded. |

***Supplementary Table 2. Journals used to identify articles for inclusion***

| **Category from InCites** | **Journal Name** | **2018 Impact Factor*** | **Abstracting & Indexing PubMed or PsychInfo?** | **Include?** |
| --- | --- | --- | --- | --- |
| “Neuroimaging” | 1. NeuroImage | 5.812 | BOTH | YES |
|  | 2. Human Brain Mapping | 4.554 | BOTH | YES |
| “Neuroscience” | 1. Nature Reviews Neuroscience | 33.162 | n/a | NO - reviews |
|  | 2. Nature Neuroscience | 21.126 | BOTH | YES |
|  | 3. Acta Neuropathologica | 18.174 | BOTH | NO - focus |
|  | 4. Behavioral and Brain Sciences | 17.194 | n/a | NO - reviews |
|  | 5. Trends in Cognitive Sciences | 16.173 | n/a | NO - reviews |
|  | 6. Journal of Pineal Research | 15.221 | NEITHER | NO – indexing |
|  | 7. Neuron | 14.403 | PubMed | YES |
| “Psychiatry” | 1. World Psychiatry | 34.024 | PsychInfo | NO - focus |
|  | 2. Lancet Psychiatry | 18.329 | BOTH | NO - focus |
|  | 3. JAMA Psychiatry | 15.916 | BOTH | NO - focus |
|  | 4. Psychotherapy & Psychosomatics | 13.744 | BOTH | NO - focus |
|  | 5. American Journal of Psychiatry | 13.655 | BOTH | YES |
|  | 6. Molecular Psychiatry | 11.973 | BOTH | YES |
| Specialty Journals (not an InCites category) | Developmental Cognitive Neuroscience | 4.920 | PubMed | YES |
|  | Journal of Cognitive Neuroscience | 3.029 | BOTH | YES |
|  | Social Cognitive and Affective Neuroscience | 3.622 | BOTH | YES |

Note. Journals were excluded due to either (1) “reviews” - journals that primarily publish review papers, (2) “focus” - journals that do not include the words “neuroimaging” or “neuroscience” in the public statement of aims and scope, and (3) “indexing” - journals that are not indexed on PubMed or PsychInfo.

***Supplementary Table 3. Codebook for full-text data extraction***

| **Question** | **Response Options** | **Exclusions and Notes for RAs** | **Reporting Score - Strict** | **Reporting Score - Loose** |
| --- | --- | --- | --- | --- |
| **Global Study Features** | | |  |  |
| Should this study be excluded from the review? If so, for what reason? | 1. Subjects are non-human  2. The imaging modality is not MRI or fMRI or there is not an imaging component of the study |  | NA | NA |
| Who are the study participants? | 1. Adults  2. Children (< 18years)  3. All ages | In the case of a longitudinal study, code this item based on the time of imaging data collection. If participants span childhood and adolescence, code as #3. | NA | NA |
| Is this a patient population? | 1. Yes  2. No | Subjects meet diagnostic criteria for a medical disorder. | NA | NA |
| What type of study is this? | 1. Observational  2. Intervention/Treatment | An intervention or treatment study must (a) randomize participants to one or more conditions, and (b) include a control group | NA | NA |
| What imaging modality was employed? | 1. functional MRI (i.e., resting state, task)  2. structural MRI (volumetric, white matter tractography, DTI, DWI)  3. Both sMRI and fMRI | We want to know what imaging modality was used in the primary analyses. Remember that sMRI images are oftentimes used to preprocess fMRI data, but this wouldn't count as a primary use of sMRI data. | NA | NA |
| How many participant samples were used in the current study? | [text entry] | This will oftentimes occur if the study replicated their results in multiple samples. For the purposes of the current review, code the primary sample or the first sample reported. | NA | NA |
| Did this study utilize data from a consortium of several participant samples? | 1. Yes  2. No | e.g., ABIDE, ENIGMA | NA | NA |
| What is the total sample size from which the imaging data was drawn from? | [text entry] | This is the largest sample size that would be reported and refers to the sample from which the imaging data was drawn. Could be the same as the imaging N, which would occur if the study was purely a neuroimaging study. | NA | NA |
| What is the largest analytic sample size with imaging data reported? | [text entry] |  | NA | NA |
| What is the smallest analytic sample size with imaging data reported? | [text entry] | In the case of group comparisons, the smallest sample size would still be the sum of both groups unless other analyses examined only one group. | NA | NA |
| **Demographic Information** | | |  |  |
| **Question** | **Response Options** | **Additional Notes for RAs** | **Reporting Score - Strict** | **Reporting Score - Loose** |
| Did the study reference a previous paper to report the demographic information of the sample? | 1. Yes  2. No |  | NA | NA |
| Which component of the study does the paper report demographic information for? | 1. All participants, whether they had imaging data or not  2. Participants with imaging data only  3. Demographics were reported for both the imaging sample and the total sample | We want demographic information about the imaging sample. However, if demographics are reported for the non-imaging component of the sample only, use that available demographic information. This question will tell us the source of the demographic information (imaging/non-imaging). | NA | NA |
| Does the study report the gender make-up of the sample (either as Ns or %) | 1. Yes  2. No |  | Yes = 1  No = 0 | Yes = 1  No = 0 |
| If gender make-up is reported, what %age of the sample is female? | [text entry] | If gender make-up is reported by group, calculate a weighted mean (using each group’s N) where possible. | NA | NA |
| Does the study report the race and/or ethnicity of the sample? | 1. Race only  2. Ethnicity only  3. Both race and ethnicity  4. Neither – there is no mention of the racial or ethnic composition of the sample  5. Something else was provided | We rely on the U.S. Census definition of Ethnicity – Hispanic/Latino or Non-Hispanic/Latino. For all of the following questions, if race or ethnicity is reported by group, calculate weighted means if possible (using each group’s N). | Race, Ethnicity, both = 1  Neither or Something else = 0 | Race, Ethnicity, both, something else = 1  Neither = 0 |
| If race is reported, what %age of the sample is reported as White? | [text entry] | Round to the nearest percent. | NA | NA |
| If race is reported, what %age of the sample is reported as Black? | [text entry] | Round to the nearest percent. | NA | NA |
| If race is reported, what %age of the sample is reported as Asian? | [text entry] | Round to the nearest percent. | NA | NA |
| If race is reported, what %age of the sample is reported as American Indian or Alaska Native? | [text entry] | Round to the nearest percent. | NA | NA |
| If race is reported, what %age of the sample is reported as Native Hawaiian or Other Pacific Islander? | [text entry] | Round to the nearest percent. | NA | NA |
| If race is reported, what %age of the sample is reported as Biracial or Multiracial? | [text entry] | Round to the nearest percent. | NA | NA |
| If ethnicity is reported, what %age of the sample is reported as Hispanic or Latino? | [text entry] | Round to the nearest percent. | NA | NA |
| If race/ethnicity is reported differently than above, list all available information here | [text entry] |  | NA | NA |
| Does the study report the mean age (in years or months) and the age range of the sample? | 1. Only age range  2. Only average age  3. Both average age and age range  4. No |  | Both average and range = 1  Only range or average = 0.5  No = 0 | Both average and range, only range, only average = 1  No = 0 |
| Does the study report the socioeconomic background of the sample? For continuous measures, mean and range. For categorical indicators, %age | 1. Income  2. Education  3. Poverty ratio  4. Some combination of 1/2/3  5. Something else [list]  6. No  7. College students | For studies where participants are children, consider family socioeconomic background. | Income, Education, Poverty Ratio, combination, something else = 1  No, college students = 0 | Income, Education, Poverty Ratio, combination, something else, college students = 1  No = 0 |
| What country is the sample from? | [text entry] | Sample country is based on where participants were recruited from. If not easily identifiable, record "Unknown". | NA | NA |
| If income is reported, is annual or monthly reported in the manuscript? | 1. Annual  2. Monthly | If subjects are minors, report primary caregiver. | NA | NA |
| If income is reported continuously, what is the average *annual* income of the sample. | [text entry] | If monthly is provided in the text, multiple by 12 for an estimate of annual income. If subjects are minors, report primary caregiver. | NA | NA |
| If income is reported categorically, what is the modal income of the sample? | [text entry] | Mode = most frequently occurring . If subjects are minors, report primary caregiver. | NA | NA |
| If income or poverty ratio is reported, what currency is reported? | [text entry] | If subjects are minors, report primary caregiver. | NA | NA |
| If education is reported categorically, what %age of the sample has a 4-year college degree or higher? | [text entry] | If subjects are minors, report primary caregiver. | NA | NA |
| If education is reported categorically, what %age of the sample has a high school degree, GED, or less? | [text entry] | If subjects are minors, report primary caregiver. Include GED completion. | NA | NA |
| If education is reported continuously, what is the average number of years of education on the sample? | [text entry] | If subjects are minors, report primary caregiver. | NA | NA |
| If poverty ratio is reported, what is the average poverty ratio of the sample? | [text entry] | Poverty ratio is calculated as the income to needs ratio, based on household income and number of people living in the home. This may be referred to as the income-to-needs ratio. | NA | NA |
| If SES is reported differently than above, list all available information here | [text entry] | Note “college students” if applicable. | NA | NA |
| **Methods Features** | | |  |  |
| **Question** | **Response Options** | **Additional Notes** | **Reporting Score - Strict** | **Reporting Score - Loose** |
| Did the study reference a previous paper to report recruitment procedures and/or data availability of the sample? | 1. Yes  2. No |  | NA | NA |
| To what extent does the study report recruitment procedures? | 1. There is explicit mention of where participants were recruited from and through what method.  2. There is explicit mention of where participants were recruited from and through what method; there were multiple recruitment methods/sites, but Ns were not broken down by method/site.  3. There is explicit mention of where participants were recruited from and through what method; there were multiple recruitment methods/sites, and Ns were broken down by method/site.  3. Some information about recruitment was stated, but details were vague  4. No information about recruitment was stated | Explicit recruitment information must include information about where participants were recruited from (i.e., location) and through what method (e.g., clinic referrals, online, posted flyers). | Explicit, including multiple sites not broken down= 1  None or some information, but details are vague = 0 | Explicit, including multiple sites not broken down = 1  Some information = 0.5  None = 0 |
| Did the study explicitly recruit cases or controls, or participants based on specific criteria? | 1. Yes  2. No | Cases/controls may refer to patient groups (e.g., having a psychiatric diagnosis) or participants based on other criteria (e.g., trauma-exposed versus non-trauma exposed). Note that if the current study created these groups as part of the analytic plan, but did not explicitly recruit separate groups, do not count this as a case/control study. | NA | NA |
| If this study recruited cases versus controls, did the study report how the recruitment method differed by case/control status? | 1. Yes, there is explicit mention of whether and how cases and controls were recruited differently  2. No, there was no mention of whether and how cases and controls were recruited differently |  | NA | NA |
| Does the study report how many participants were initially contacted and how many agreed to participate in the study versus declined? | 1. Yes, the study reported the number of participants who were initially contacted, N who agreed, and N who refused to participate  2. No, the study did not report any information about initial recruitment efforts |  | Yes = 1  No = 0 | Yes = 1  No = 0 |
| Does the study report a date range for the overall data collection? | 1. Yes  2. No | Imaging or non-imaging data | Yes = 1  No = 0 | Yes = 1  No = 0 |
| Does the study report clear inclusion and exclusion criteria for participation in the study? | 1. Yes, inclusion/exclusion criteria were explicitly stated.  2. Some information about inclusion/exclusion criteria were stated, but details were vague  3. No reference to inclusion/exclusion criteria | To qualify as a “1” for this question, studies must explicitly reference 1 or more inclusion/exclusion or eligibility/ineligibility criteria in the text. If the study used language such as “all participants were free of psychiatric disorders”, code this as #2. | Yes = 1  Some but vague, No = 0 | Yes = 1  Some but vague = 0.5  No = 0 |
| Does the study report the number of **participants** (if any) refused or were unable to complete the neuroimaging protocol? | 1. Yes, the study provided a total number and listed reasons why there was no neuroimaging data  2. Yes, the study provided a total number, but did not list reasons why there was no neuroimaging data.  3. Yes, the study provided the number of participants with no neuroimaging data for each of several categories (e.g., N=X refused, N=X braces).  4. No, there was no information about the number of participants who were part of the overall study but who did not have neuroimaging data. | Examples of reasons for no neuroimaging data could include (but are not limited to) refusal, metal in body, medical reasons. Note that if participation in the neuroimaging study was an inclusion criterion and every participant completed the scan, then this question should be coded as “1”. | Yes with total number and reasons, Yes with total number and reasons in specific categories = 1  Yes with a total number but no reasons, No information = 0 | Yes with total number and reasons, Yes with total number and reasons in specific categories = 1  Yes with a total number but no reasons = 0.5  No information = 0 |
| Does the study report the number of participants (if any) were removed from the analysis due to poor data quality? | 1. Yes, the study provided a total number and listed reasons for poor data quality  2. Yes, the study provided a total number, but did not list reasons for poor data quality.  3. Yes, the study provided the number of participants removed for poor data quality for each of several categories (e.g., N=X movement, N=X warping).  4. No, there was no information about the number of participants who were removed from the analyses due to poor neuroimaging data quality | Examples of reasons for poor data quality include (but are not limited to) movement, ghosting or warping of images, low accuracy, falling asleep. If no participants were excluded due to poor data quality and the article does not explicitly state this, code as #4. If the article explicitly states that all participants passed data quality checks, code as #1, #2, or #3 depending on the detail of information provided. | Yes with total number and reasons, Yes with total number and reasons in specific categories = 1  Yes with a total number but no reasons, No information = 0 | Yes with total number and reasons, Yes with total number and reasons in specific categories = 1  Yes with a total number but no reasons = 0.5  No information = 0 |
| Was a missing values analysis completed, comparing participants with and without imaging data on demographic or study variables? | 1. Yes  2. No | These analyses might compare groups by gender, race, SES, age, or other study-relevant variables (e.g., psychopathology, cognition). This type of analysis is intended to demonstrate if there are systematic differences in who had usable imaging data (and thus were included in the analyses) and who did not. | Yes = 1  No = 0 | Yes = 1  No = 0 |
| **Generalizability and Open Science Features** | | | | |
| **Question** | **Response Options** | **Additional Notes** | **Reporting Score - Strict** | **Reporting Score - Loose** |
| Was there any mention of a target population (i.e., what population the study is trying to generalize to)? | 1. Yes  2. No | This must be an explicit generalization to a broader population. The text may include “target population”, mention that the results generalize to a broader population, or provide comparisons of estimates to population-based samples. | Yes = 1  No = 0 | Yes = 1  No = 0 |
| In the limitations section of the paper, do the authors comment on how generalizable their results are to a broader population? | 1. Yes  2. No | Authors must explicitly refer to the demographics or characteristics of the study sample as a reason for limited generalizability. | Yes = 1  No = 0 | Yes = 1  No = 0 |
| Did the paper include a pre-study power analysis? | 1. Yes  2. No  3. No, but the study presented post-hoc power analyses | Any power analysis must include notation of an effect size, statistical power, and a significance level. | Yes = 1  No = 0 | Yes = 1  No = 0 |
| Was any part of the analyses pre-registered? | 1. Yes  2. No |  | Yes = 1  No = 0 | Yes = 1  No = 0 |

***Supplemental Table 4. Study Characteristics***

| Characteristic | | N (%) |
| --- | --- | --- |
| Journal | |  |
|  | *NeuroImage* | 393 (43.0%) |
|  | *Human Brain Mapping* | 260 (28.0%) |
|  | *Social Cognitive and Affective Neuroscience* | 75 (8.2%) |
|  | *Developmental Cognitive Neuroscience* | 61 (6.6%) |
|  | *Journal of Cognitive Neuroscience* | 55 (6.0%) |
|  | *Molecular Psychiatry* | 32 (3.5%) |
|  | *The American Journal of Psychiatry* | 20 (2.2%) |
|  | *Nature Neuroscience* | 16 (1.7%) |
|  | *Neuron* | 7 (0.8%) |
| Sample Age Group | |  |
|  | Adults | 680 (74.0%) |
|  | Children (< 18 year) | 140 (15.2%) |
|  | All Ages | 99 (10.8%) |
| Patient Sample (i.e., meet diagnostic criteria for a medical disorder) | |  |
|  | No | 691 (75.2%) |
|  | Yes | 228 (24.8%) |
| Consortia-level Data Used | |  |
|  | Yes | 119 (12.9%) |
|  | No | 800 (87.1%) |
| Study Type | |  |
|  | Observational | 878 (95.5%) |
|  | Treatment / Intervention | 41 (4.5%) |
| Imaging Modality | |  |
|  | Structural MRI (e.g., volumetric, white matter tractography) | 226 (24.6%) |
|  | Functional MRI (i.e., resting-state, task-based) | 624 (67.9%) |
|  | Both Structural and Functional MRI | 69 (7.5%) |

**Supplemental Table 4. Study Characteristics are Associated with Reporting Frequency, Using Loose Reporting Criteria**

|  |  | **Demographics** | **Methods** | **Open Science & Generalizability** | **Overall Reporting Index** |
| --- | --- | --- | --- | --- | --- |
|  |  | **B (SE), β** | **B (SE), β** | **B (SE), β** | **B (SE), β** |
| **Sample Size** | | 0.000034 (0.00005), 0.02 | -0.00006  (0.00008), -0.03 | 0.00007 (0.00005), 0.05 | 0.000046 (0.00013), 0.01 |
| **Patient Population** | | 0.04 (0.06), 0.02 | 0.20 (0.10), 0.07* | 0.16 (0.06), 0.10** | 0.39(0.15), 0.09* |
| **Consortia Data** | | -0.27(0.08), -0.12*** | -0.30 (0.12), -0.08* | 0.10 (0.07), 0.05 | -0.48(0.19), -0.08* |
| **Treatment / Intervention** | | -0.04 (0.12), -0.009 | 0.59 (0.19), 0.10** | 0.09 (0.11), 0.03 | 0.64 (0.29), 0.07* |
| **Sample Age** | | F(2, 40.61) = 3.53*** | F(2, 37.50) = 14.12*** | F(2, 4.61) =  4.97** | F(2,92.08) = 14.33*** |
|  | Children | (ref) | (ref) | (ref) | (ref) |
|  | Adults | -0.16 (0.08),  -0.09* | -0.62 (0.12),  -0.22*** | -0.21 (0.07),  -0.13** | -0.98 (0.19),  -0.23*** |
|  | All ages | -0.21 (0.10), -0.08* | -0.26 (0.15), -0.07 | -0.06 (0.09), -0.03 | -0.53 (0.24),-0.09* |
| **Modality** | | F(2,19.83) = 17.87*** | F(2,2.60) =  1.00 | F(2,5.64) =  6.08** | F(2,54.71) = 8.52*** |
|  | Structural MRI | (ref) | (ref) | (ref) | (ref) |
|  | Functional MRI | -0.30 (0.06),  -0.18*** | -0.12 (0.10),  -0.05 | -0.20 (0.06),  -0.13*** | -0.62 (0.15),  -0.15*** |
|  | Multimodal | -0.24 (0.10),  -0.08* | 0.01 (0.16),  0.003 | -0.10 (0.10),  -0.04 | -0.33 (0.25),  -0.04 |
| **Journal** | | F(8,13.58) =  3.08*** | F(8,44.53) = 4.19*** | F(2,4.31) =  1.16 | F(8,101.31) = 3.94*** |
|  | *NeuroImage* | (ref) | (ref) | (ref) | (ref) |
|  | *DCN* | 0.06 (0.11), 0.02 | 0.56 (0.18), 0.11** | -0.05 (0.10), -0.02 | 0.58 (0.27), 0.08* |
|  | *HBM* | 0.18 (0.06), 0.10** | 0.20 (0.10), 0.07* | -0.06 (0.06), -0.04 | 0.32 (0.15), 0.07* |
|  | *JOCN* | -0.10 (0.11), -0.03 | 0.17 (0.17), 0.03 | -0.08 (0.10), -0.03 | -0.10(0.26), -0.002 |
|  | *Mol Psych* | 0.14 (0.14), 0.03 | 0.56 (0.22), 0.08* | 0.17 (0.13), 0.04 | 0.87 (0.35), 0.08* |
|  | *Nature Neuro* | -0.10 (0.19), -0.02 | 0.98 (0.30), 0.11*** | 0.15 (0.18), 0.003 | 0.90 (0.46), 0.06 |
|  | *Neuron* | -0.31 (0.28), -0.04 | -0.35 (0.44), -0.03 | -0.35 (0.26), -0.04 | -1.02 (0.69), -0.05 |
|  | *SCAN* | 0.30 (0.09), 0.11** | 0.36 (0.15), 0.08* | 0.13 (0.09), 0.05 | 0.80(0.23), 0.11*** |
|  | *AJP* | 0.42 (0.18), 0.08* | 0.88 (0.28), 0.11** | 0.05 (0.16), 0.009 | 1.35 (0.43), 0.10** |

*Note.* N = 919; Sample size refers to the largest sample size reported, which was first was winsorized to +3SD from the mean, and log-transformed. Multimodal refers to studies that included both structural and functional MRI. One-way ANOVAs for categorical variables evaluate the change in model fit when that predictor was removed from the model, compared to the full model. The reporting indices here reflect “loose” definitions; * *p* < 0.05, ** *p* < 0.01, *** *p* < 0.001.

***Supplementary Figure 1. Demographics, Methods, and Generalizability & Open Science Reporting using Loose Definitions***

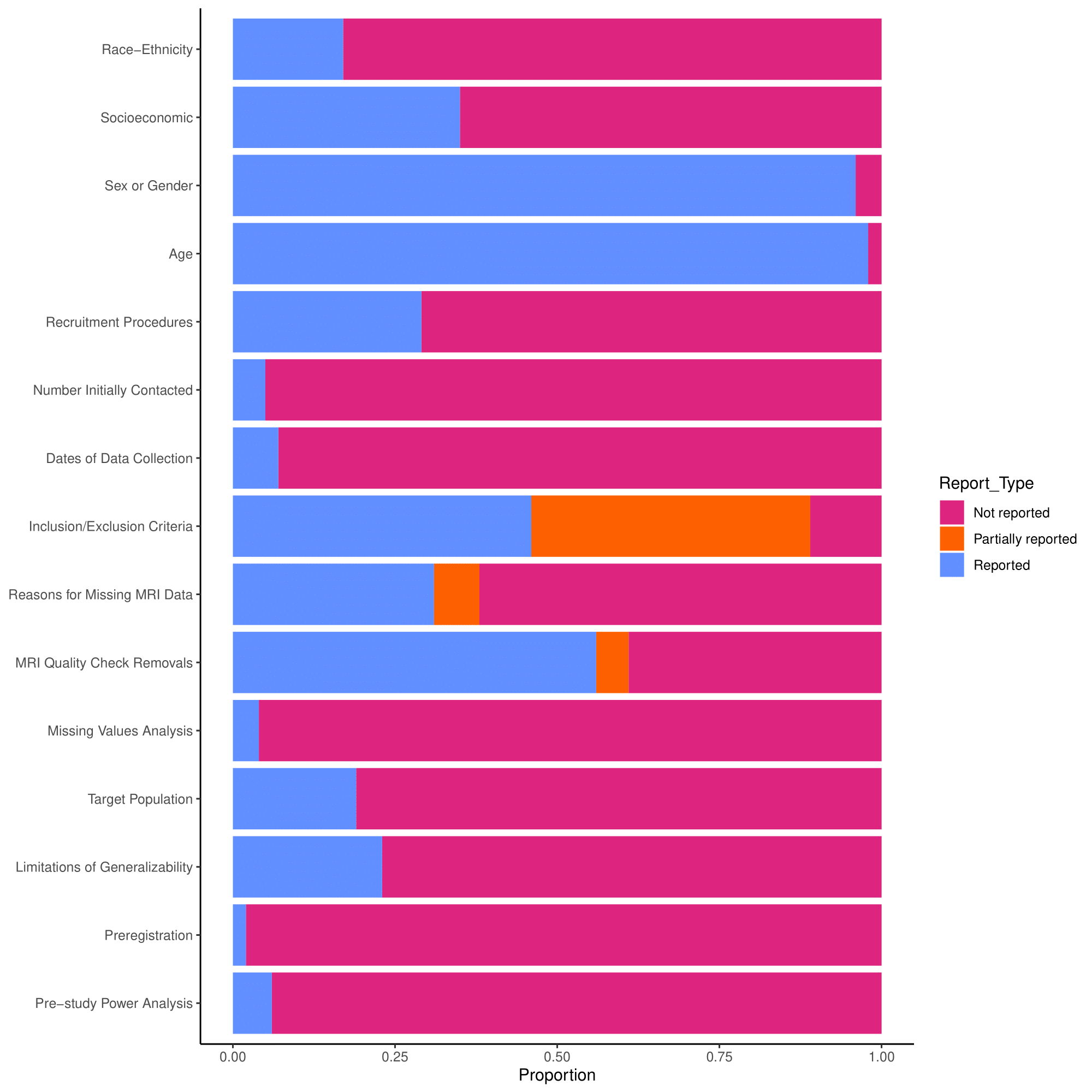

Note. N = 919 records. Definitions of whether a feature was reported, partially reported, or not reported can be found in Supplementary Table 2.
